# Distinct malignant cell states and myeloid glutamate signaling associated with aggressive pancreatic neuroendocrine tumors

**DOI:** 10.1101/2025.07.15.664730

**Authors:** Jeanna M. Arbesfeld-Qiu, Jae-Won Cho, Phuong T.T. Nguyen, Nicole A. Lester, Jennifer Su, Jimmy A. Guo, Hannah Hoffman, Carina Shiau, Nicholas Caldwell, Shugo Muratani, Miranda Galvan, Jessica E. Proctor, Zack Ely, Steven Wang, Maria Ganci, Ruben Dries, Theodore Hong, Jennifer Wo, Genevieve Boland, Carlos Fernandez-del Castillo, Cristina Ferrone, Christopher M. Heaphy, M. Lisa Zhang, Mari Mino-Kenudson, Martin Hemberg, William L. Hwang

## Abstract

Pancreatic neuroendocrine tumors (PNET) are rare malignancies of the endocrine pancreas with diverse clinical outcomes. While some PNETs are indolent, others are aggressive and metastasize quickly. However, clinically-relevant molecular stratification for PNET to predict outcomes and guide therapeutic decision-making is limited. Thus, there is an urgent need to understand the molecular heterogeneity of PNETs to refine prognostication and discover novel therapeutic vulnerabilities. We performed single-nucleus RNA sequencing on resected primary and metastatic PNETs (*n* = 20), including two PNETs with neoadjuvant treatment. We inferred gene expression programs (GEPs) of malignant and non-malignant cells and investigated associations with clinical outcomes. Next, we inferred interactions in the tumor microenvironment (TME) and performed transwell assays for functional validation. Finally, we explored genomic and transcriptomic evolution in a unique case study of an untreated primary PNET with two asynchronous hepatic metastases. A malignant GEP enriched for neural/synaptic signaling genes was associated with worse overall survival, broad chromosomal loss of heterozygosity, and alternative lengthening of telomeres. Another malignant GEP enriched for VEGF signaling increased throughout metastatic progression in our case study. We found that macrophage-derived glutamate drives polarization towards an immunosuppressive phenotype and activates the MAPK/ERK pathway in malignant cells to increase migratory capacity. This study provides a detailed single-nucleus transcriptomic classification of malignant, stromal, and immune cell types and states in PNETs, their interactions in the TME, and associations with clinical outcomes. The refined molecular taxonomy of PNET may guide the development of more efficacious biomarkers and therapeutic strategies.

## Introduction

Pancreatic neuroendocrine tumors (PNETs) derive from the neuroendocrine cells in the pancreas, with an average incidence of 1 per 100,000 individuals.^1^ PNETs are categorized as “functional” (10%) or “non-functional” (90%) based on production or absence of excess hormones, respectively.^1^ Since non-functional PNETs often lack symptoms until a significant tumor burden is reached, they are typically found at advanced stages. While early-stage disease has a five-year survival rate of 95%, about 40-80% of patients ultimately develop metastases, with five-year survival dropping to 25% in the case of distant metastases.^2^ There are limited treatment options for patients with progressive PNET, including somatostatin analogues, mTOR pathway inhibitors, and multi-kinase inhibitors.^1^ However, therapeutic selection is not currently driven by the molecular features of the tumor and treatment resistance is pervasive.^3^ A greater understanding of the biological pathways critical for malignant neuroendocrine cell survival and proliferation is necessary to identify new therapeutic vulnerabilities in PNET.

Our understanding of PNET tumorigenesis and progression is limited by the lack of genetically-tractable *in vitro* and *in vivo* models that recapitulate the human disease.^4,5^ Current models are limited in that most represent functional PNETs or more aggressive neuroendocrine carcinomas, which constitute the minority of PNETs. An alternative approach has been molecular profiling of patient tumors to gain insights into PNET biology, but such efforts have been limited by the paucity of patient samples. Nevertheless, genomic profiling of patient tumors has revealed that the most common pathways altered in PNET include chromatin modification (*MEN1*, *SETD2, MLL3*), alternative lengthening of telomeres (*DAXX, ATRX)*, DNA damage repair (*BRCA2, CHEK2, MUTYH)*, and mTOR signaling (*PTEN, TSC1/2*).^6,7^ Furthermore, previous studies using bulk molecular profiling have revealed some clinically-relevant subgroups:^8–14^ beta cell-like state driven by the transcription factor PDX1 that is associated with low copy number burden and better prognosis;^8,12,14^ an alpha cell-like state driven by variable ARX activity, mutations in *ATRX*, *DAXX*, or *MEN1,* and loss of chromosome 11 heterozygosity;^10,14,15^ as well as tumors with predominant stromal/mesenchymal or proliferative features.^12^ Furthermore, subtypes associated with high metastatic potential and poor prognosis tend to be associated with genes related to early pancreas development, hypoxia, and inflammatory pathways;^8,13^ high copy number burden; and broad loss of heterozygosity in a set of ten chromosomes (chromosomes 1, 2, 3, 6, 8, 10, 11, 16, 21, and 22).^9,10,16^ Although these studies provide a starting point for unraveling the molecular determinants of PNET biology and clinical outcomes, an important limitation is that they studied entire tumor samples in aggregate, resulting in an unknown mixture of different cell types and states and rendering it difficult to distinguish specific features of malignant cells from other cells in the microenvironment.

Our understanding of how cancer cell-extrinsic features in the tumor microenvironment (TME) contribute to disease progression in PNETs is limited. However, it has become clear that myeloid cells contribute to poor clinical outcomes. For example, immunohistochemical analysis of human PNET samples have shown that tumor associated macrophage (TAM) infiltration is associated with worse prognosis^17–19^ and deficiency of CSF-1 disrupts PNET tumorigenesis.^20^ Few studies have explored the cell state of TAMs in PNET, although it was recently shown that CD163-high macrophage abundance was correlated with worse prognosis.^18^ Moreover, a small single-cell RNA-seq analysis of gastroenteropancreatic neuroendocrine tumors found that myeloid cells express immunosuppressive ligands.^21^

Single-cell/single-nucleus RNA-seq enables detailed characterization of individual malignant and non-malignant cells within a tumor. In particular, single-nucleus RNA-seq (snRNA-seq) has the benefits of being able to profile frozen, archival specimens, often with better representation of cell type distribution in tissues that are difficult to dissociate.^22,23^ While previous studies have applied single-cell RNA-seq to patient-derived PNET samples to highlight the heterogeneity of cells in the TME, a detailed investigation into the diversity of malignant cell states and their interactions with other cell types in the TME has yet to be performed at scale.^21,24,25^

In this study, we used snRNA-seq to profile a cohort of 20 surgically resected PNET specimens, recovering 125,681 high quality single-nucleus profiles. We assembled a *de novo* taxonomy of malignant cell states across the subset of non-functional PNETs (*n =* 14), including two novel cell states that are enriched in neuronal/synaptic signaling genes and VEGF-signaling genes, respectively. We also characterized nonmalignant cell states across cancer-associated fibroblasts, vascular endothelial cells, and macrophages. By inferring cell-cell interactions from our snRNA-seq profiles, we predicted paracrine interactions between malignant cells and other cell types within the TME. Finally, in a unique case study of a primary PNET with two asynchronous metastases, we show that the VEGF-signaling malignant subtype is increased throughout progression and interacts with immunosuppressive macrophages via glutamate, suggesting these pathways and interactions as potential targets for therapeutic intervention.

## Results

### Characterizing the tumor microenvironment of primary untreated pancreatic neuroendocrine tumors at single-cell resolution

We performed snRNA-seq on a cohort of 20 patient-derived PNET specimens (**Fig. 1a**; **Table S1**; **Methods**). This cohort included primary PNETs (n = 18) and hepatic metastases (n = 2). Most tumors were non-functional (n = 19), except for one glucagonoma. This cohort encompassed all tumor grades, and most samples were untreated (n = 18) with two samples having received neoadjuvant treatment. After quality control filtering, we recovered 125,681 high-quality single nucleus profiles.

**Figure 1.**
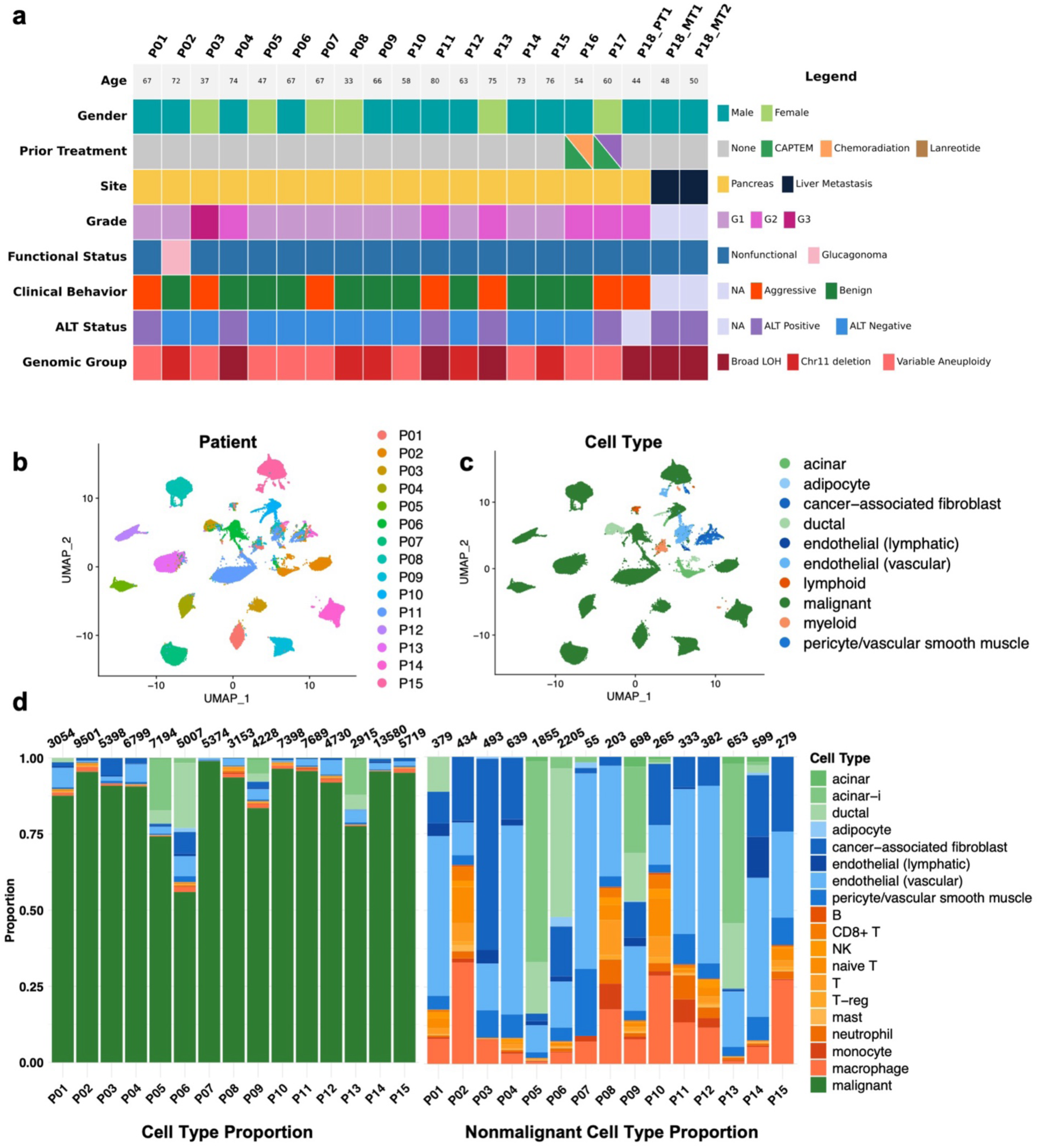
Single-nucleus RNA-sequencing of primary, untreated pancreatic neuroendocrine tumors captures diversity of cell types. **(a)** Clinical information of PNETs profiled with snRNA-seq**. (b,c)** Uniform manifold approximation and projection (UMAP) embedding of single-nucleus profiles (dots) of all cells from the 15 primary, untreated PNET tumors, colored by patient ID (**b**, legend) and cell type (**c**, legend). **(d)** Stacked barplot of cell type proportion of each specimen with (left) and without (right) malignant cells. Number of cells per sample is indicated above each bar.

First, we investigated the cellular landscape of untreated primary PNETs to dissect the composition of the TME without treatment perturbation (n = 15 specimens; 91,869 nuclei). We identified malignant cells by expression of canonical PNET marker genes (e.g., *SYP, CHGA, CHGB, NCAM1*) and confirmed they had a higher copy number burden compared to nonmalignant cells (**Fig. S1**; **Methods**). The malignant cells formed separate clusters by patient (**Fig. 1b**). Next, we used unsupervised clustering and marker genes to identify major cell types among the nonmalignant cells (**Fig. 1c**). We subclustered each non-malignant cell type and used established gene signatures to annotate detailed cell types (**Fig. 1c-d**, **Fig. S2**).^26,27^ We identified various nonmalignant cell types including diverse stromal cells (cancer-associated fibroblasts, endothelial cells, pericytes/vascular smooth muscle, and adipocytes), nonmalignant epithelial cells (ductal, acinar), and a variety of immune cells. The most abundant acinar cell subtype was an acinar-idling cell, a subtype thought to have reduced protein secretion and increased response to external stimuli from islets.^27^ Overall, there were more myeloid than lymphoid cells, particularly macrophages, consistent with findings from prior single-cell transcriptomic studies of GEP-NETs.^21,25^

### Malignant gene expression programs shared across non-functional PNETs

Next, we investigated malignant cell states in treatment-naive primary non-functional PNETs which are the most common clinical subtype of PNET. Using the 73,200 malignant cells derived from these tumors, we used consensus non-negative matrix factorization (cNMF) as an unbiased approach to infer gene expression programs (GEPs) (**Fig. 2a**). Using established procedures (**Methods**), we selected 25 GEPs (**Fig. S3a**) and focused on the 23 GEPs that were robust to downsampling (**Fig. S3b**; **Methods**). Of these 23 GEPs, 16 programs were patient-specific, and we did not analyze them further as we were primarily interested in investigating GEPs that were broadly applicable to PNET biology (**Fig. S3c**). We annotated the remaining seven GEPs by their top 100 weighted genes, compared them to published bulk-derived signatures of PNETs^12^ and physiological neuroendocrine cell subtypes,^26^ and performed gene set overrepresentation analysis (**Fig. S3d, S4a Table S2**).^28^ Two GEPs had significant overlap with known signatures: Malig-5 with the beta-cell-like signature (hypergeometric test adjusted p-value = 1.47×10^−33^), and Malig-7 with the proliferative signature (hypergeometric test adjusted p-value = 2.03×10^−105^). Malig-1 and Malig-2 were enriched for ribosomal and mitochondrial genes, respectively. The remaining three GEPs without overlap with known signatures included Malig-3, which was enriched for genes related to neural/synaptic signaling (*NRXN3, NTNG1, GRIA1*); Malig-4, which was enriched for genes related to VEGF-signaling (*NR4A1/3, EGR3, ITGAV*); and Malig-6, which was enriched for genes related to inflammation and interferon-gamma signaling (**Fig. 2b, S4a**).

**Figure 2.**
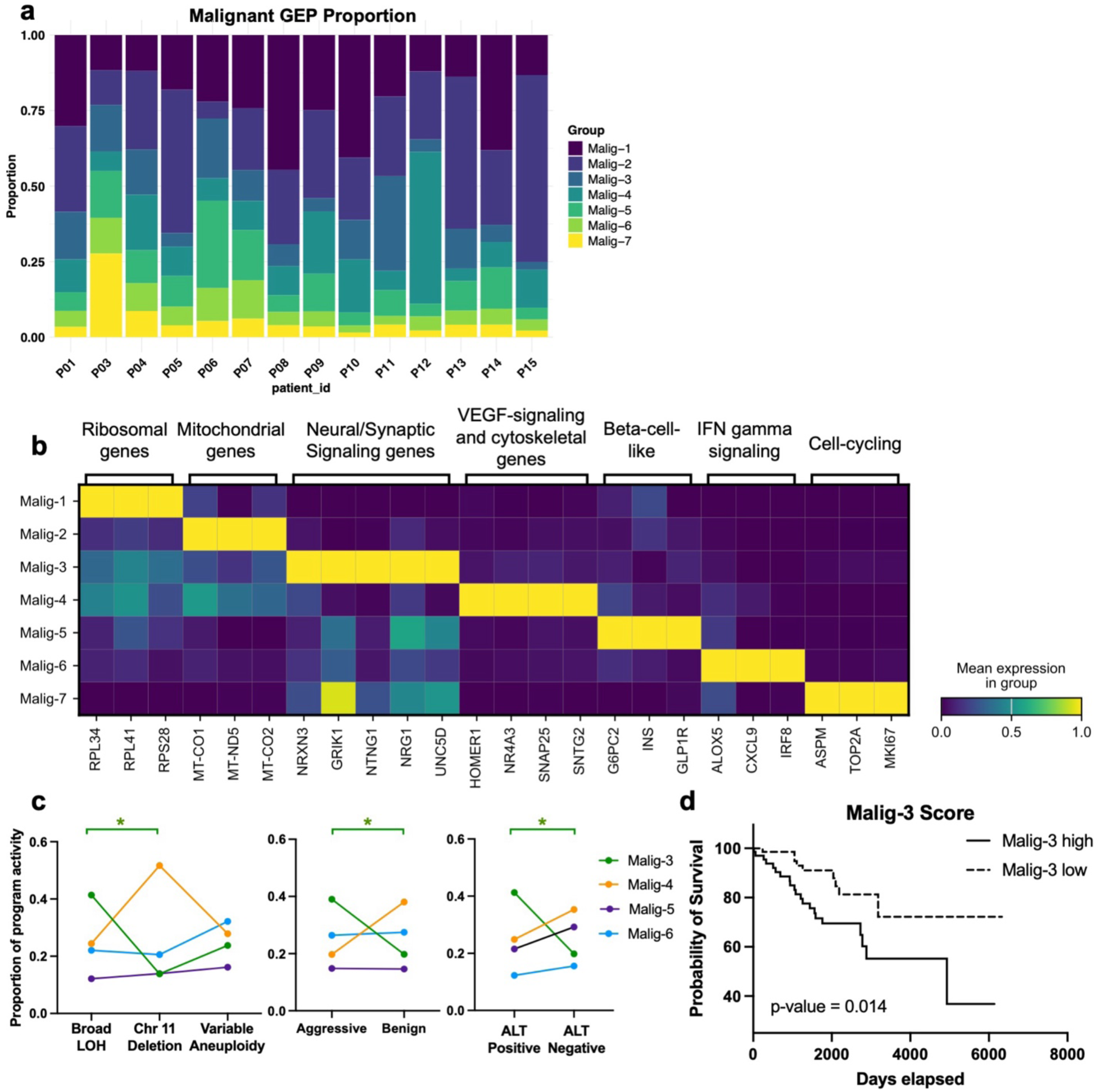
Consensus NMF recapitulates known bulk malignant programs and identifies two novel gene expression programs (GEP) in nonfunctional PNETs. **(a)** Distribution of GEP activity (legend) across each nonfunctional PNET in our cohort (columns). **(b)** Matrix plot of scaled average expression of top-weighted genes in each cNMF program with representative gene family terms for each GEP indicated (top). **(c)** Proportion of GEP activity (legend) across genomic subgroups of PNET (left) and aggressive versus benign clinical activity (middle) and ALT positive versus negative (right) in our primary cohort. Statistical significance was determined by FDR < 0.1 in a linear mixed model with patient ID as a random effect. Color of asterisk indicates the GEP with significant activity (e.g. green – Malig-3). **(d)** Kaplan Meier plot of overall survival (days) between Malig-3 high samples and Malig-3 low samples from external bulk RNA cohorts.

To identify potential transcription factor (TF) regulators of these cell states, we performed single-cell regulatory network inference and clustering (SCENIC) on the malignant single-nucleus profiles from the non-functional PNET cohort.^29^ We identified TFs whose activity was significantly correlated with GEP scores in malignant cells (**Fig. S4b**). Consistent with previous findings, the TFs with highest correlation with Malig-5, which resembled a beta cell-like state, were *PDX1* and *NKX6-2*, two TFs associated with promoting beta-cell fate in the developing pancreas.^30^ The TFs with highest correlation with Malig-7, which was enriched for cell-cycling genes, included *E2F* family TFs, which regulate genes required for progression through the cell cycle, and have pro-oncogenic effects in various malignancies.^31^ Other TFs significantly associated with Malig-7 included known oncogenes (*MXD3*, *MYBL1)* that mediate progression through the cell cycle.^32,33^ Malig-3 had the highest correlation with *TCF12*, a TF associated with progression and metastasis in colorectal cancer, glioblastoma, and gallbladder cancer,^34^ and *SOX4*, a TF involved in stemness and neuronal differentiation, and associated with epithelial to mesenchymal transition in multiple cancers.^35^ Malig-4 had the highest correlation with TFs implicated in VEGF-signaling and angiogenesis, such as FOS family members (*FOSB, FOSL2*),^36,37^ as well as *ATF6*, which was also one of the top weighted genes in this GEP.^38^ Malig-6 had highest correlation with *BNC2*, a transcription factor that upregulates interferon-stimulated genes in lung cancer models and is implicated in increased invasion and epithelial-mesenchymal transition in other cancers.^39–41^

### Malig-3 GEP is associated with loss of heterozygosity and worse overall survival

Next, we investigated if these malignant cell states correlate with previously reported genomic subtypes of PNET.^9,10^ Using inferred copy number variations, we categorized the non-functional PNETs in our cohort into previously described genomic subtypes: Group 1 with loss of heterozygosity in chromosomes 1, 2, 3, 6, 8, 10, 11, 16, 21, 22 and associated with poor clinical outcome; Group 2 with loss of heterozygosity of chromosome 11 and associated with intermediate clinical outcome; and Group 3 with variable aneuploidy, associated with better clinical outcome (**Fig. S1; Table S1; Methods**). Of note, although we did not include the glucagonoma P02 in this analysis, it featured a deletion of chromosome 11, which is consistent with previous studies showing Group 2 PNETs being associated with *GCG* expression (**Fig. 1a**).^9^ Group 1 was represented in 21.4% of our cohort (3/14), 28.5% (4/14) were in Group 2, and 50% (7/14) were in Group 3. Next, we calculated the proportion of GEP activity in each of these genomic groups (**Fig. 2c**). As the inclusion of ribosomal (Malig-1), mitochondrial (Malig-2), and cell-cycling (Malig-7) genes would be less informative for specific features of PNETs, we focused on the activity of Malig-3, Malig-4, Malig-5, and Malig-6. Malig-3 had the highest activity in Group 1 tumors (FDR = 0.003; Gamma mixed effects regression model, **Methods**). The Malig-4 GEP trended towards greater activity in Group 2 tumors. To investigate if the GEPs identified had an association with clinical outcomes in our cohort, we divided our cohort into tumors with aggressive behavior (n = 5), defined as G3, nodal metastases, or ultimate development of distant metastasis, and all others as benign behavior (n = 9). Again, we found that Malig-3 was significantly enriched in PNETs with aggressive behavior (**Fig. 2c**; FDR = 0.026, Gamma mixed effects regression model). Next, we investigated if the GEPs had an association with ALT, as this is a prognostic biomarker in PNET.^42–44^ Malig-3 was significantly associated with ALT-positive status (FDR = 0.043, Gamma mixed effects regression model). Conversely, there was a trend towards enrichment of Malig-4 and Malig-5 in PNETs with ALT-negative status (**Fig. 2c**).

To examine the generalizability of Malig-3, Malig-4, Malig-5, and Malig-6 for prognostication, we scored clinically-annotated bulk RNA-seq data of primary PNETs from three previously published cohorts with a combined total of 140 cases.^11,12,15^ We found that high Malig-3 score was significantly associated with worse overall survival (log-rank p-value = 0.014; **Fig. 2d**), while Malig-4, Malig-5, and Malig-6 did not stratify overall survival (**Fig. S4c**). Together, these results show that Malig-3, the GEP featuring neural/synaptic signaling genes, is associated with worse prognosis in several independent cohorts of primary PNETs.

### Neoadjuvant treatment is associated with diverse cancer cell states

Treatment is known to modulate cancer cell state and TME composition.^45^ To explore the effects of treatment on PNETs, we analyzed two primary grade 2 PNETs (P16 and P17) that received neoadjuvant therapy in the form of capecitabine and temozolomide (CAPTEM) with (P16) or without additional chemoradiation (P17). In follow-up, the patient that received CAPTEM alone (P17) progressed to having liver metastases while the patient that received CAPTEM and chemoradiation (P16) had no evidence of disease (**Fig. 1a; Table S1**).

After cell annotation (**Fig. S5a-b**), we scored each malignant cell for the GEPs identified from the untreated cohort (**Fig. S5c**). We found that P16 had the highest score of Malig-5, the beta cell-like GEP, consistent with the more favorable clinical course for this patient. In contrast, P17 was not clearly enriched for any of the GEPs identified in the untreated cohort. When examining the differentially expressed genes in the malignant cells from P17 compared to the malignant cells from P16, the highest fold-change genes included those typically derived from the acinar lineage, such as *PTF1A*, a transcription factor associated with the maintenance of pancreatic exocrine function,^46^ and various digestive enzymes (*PRSS1, CTRB2, CELA3A*) (**Fig. S5d**). Although a small sample size, these results suggest that the malignant cell states in treated primary PNETs are not fully captured by the GEPs identified in untreated primary PNET.

### Nonmalignant cell states across non-functional PNETs

As snRNA-seq allowed us to investigate all cells in the TME at single-cell resolution, we also inferred GEPs in key nonmalignant cell types across the treatment-naive non-functional primary PNETs in our cohort (n = 14) (**Table S3)**. While previous studies have used conventional unsupervised clustering to describe heterogeneity in nonmalignant cell types in PNET, we used topic modelling with cNMF as this method captures continuous GEPs shared by cells, rather than forcing discrete clusters. Below, we highlight some of the key programs identified.

#### Cancer-associated fibroblasts (CAFs)

Cancer-associated fibroblasts (CAFs) have been shown to play an important role in tumorigenesis and disease progression in other cancers.^47^ Thus, we subclustered the 1,367 single-nucleus profiles annotated as CAFs and cNMF revealed three GEPs (**Fig. 3a-b, S6a**). Previously identified signatures for CAF subtypes did not align well with the CAF subsets in PNET (**Fig. S6b**).^48,49^

**Figure 3.**
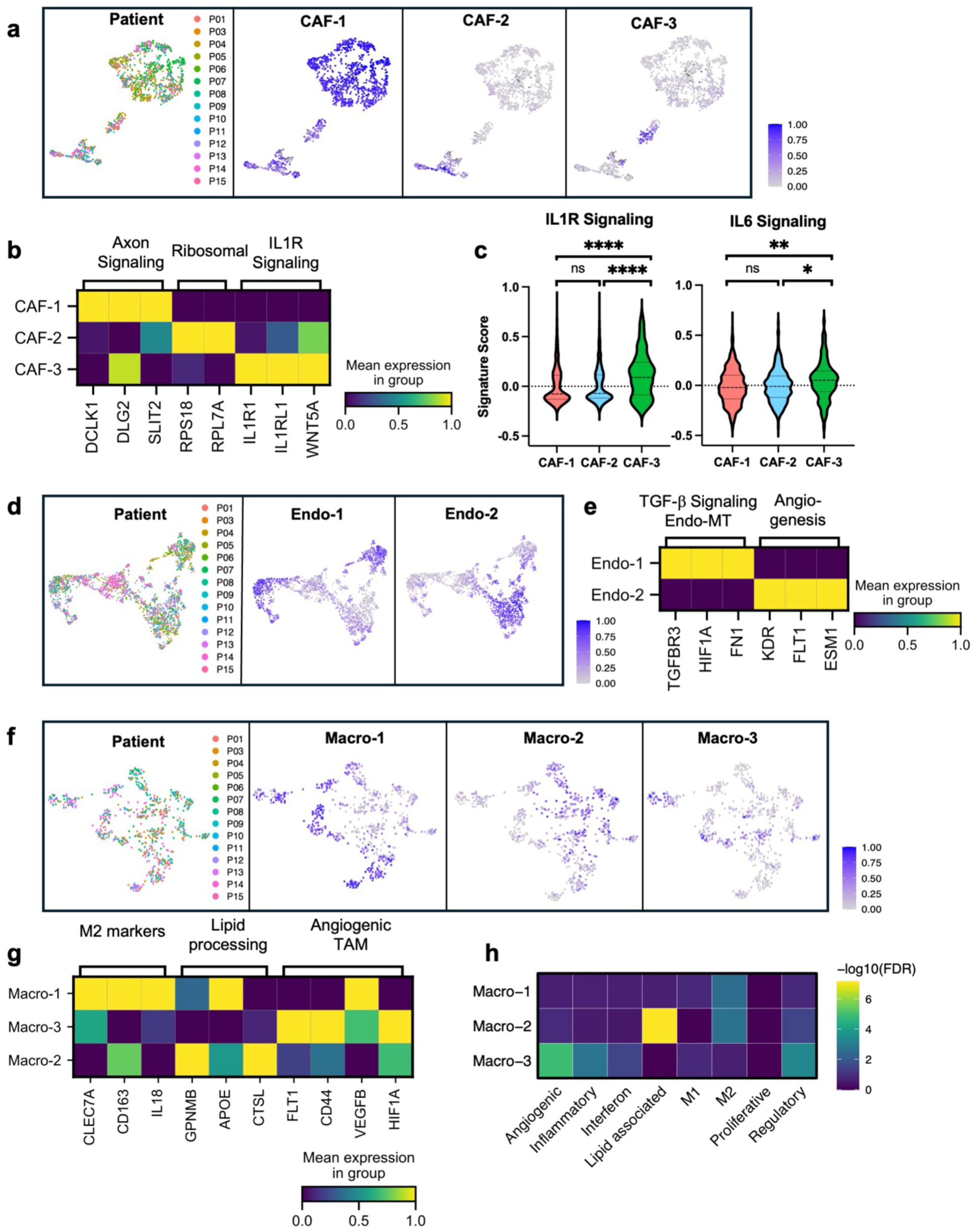
Nonmalignant cell states across cancer-associated fibroblasts, vascular endothelial cells, and macrophages in primary, nonfunctional PNET. **(a)** UMAP embedding of single-nucleus profiles (dots) of fibroblasts in our primary cohort colored by sample (left) and the normalized expression score (blue) of each fibroblast GEP. **(b)** Matrixplot of average scaled gene expression of top-weighted CAF GEP genes with representative gene family terms labeled (top). **(c)** Barplots comparing score of Gene Ontology “Molecular Function Interleukin-1 Receptor Activity” gene set (top) across fibroblast subtypes (columns) and the Gene Ontology “Biological Processes Interleukin-6 Signaling” gene set (bottom) across fibroblast subtypes. Asterisks represent significance of adjusted p-value with Holm’s from pairwise Wilcoxon t-test: **** p <0.0001, *** p < 0.001, ** p < 0.01, * p < 0.05, ns p ≥0.05 **(d)** UMAP embedding of single-nucleus profiles (dots) of vascular endothelial cells in our primary cohort colored by sample (left) and the normalized expression score (blue) of each vascular endothelial GEP. **(e)** Matrixplot of average scaled gene expression of top-weighted endothelial GEP genes with representative gene family terms labeled **(f)** UMAP embedding of single nucleus profiles (dots) of macrophages from primary, nonfunctional, untreated PNET tumors colored by the normalized expression score (blue) of each macrophage GEP. **(g)** Matrixplot of average scaled gene expression of top-weighted CAF GEP genes with representative gene family terms labeled (top). **(h)** Similarity of macrophage GEPs (labels, rows) compared to single-cell signatures of macrophage subtypes (columns). Statistical significance of overlap (-log10(adjusted p-value), hypergeometric test) for each pair of de novo cNMF annotated program and prior signature indicated by color bar. Pairs with no overlap are shown in black.

CAF-1 corresponded to the largest subcluster of CAFs and was enriched for gene ontology terms related to nervous system development (e.g. *DCLK2*, *DLG2*, *NAV3*) and axon-guidance (*SLIT2/3*) (**Fig 3b, S6c**). CAF-1 also had significant overlap with the “neurotropic CAF” GEP identified in PDAC (hypergeometric test p-value = 3.50 × 10^−10^).^22^ CAF-2 corresponded to the next largest cluster, and was enriched for ribosomal genes, suggesting higher protein synthesis rates (**Fig 3b, S6c**).^50^ Finally, CAF-3 was enriched for genes related to IL1R signaling and had significantly higher scores of downstream IL1R and IL6 signaling compared to CAF-1 and CAF-2 (**Fig. 3b-c**), suggesting a more inflammatory phenotype. Although there was no significant overlap between this GEP and the iCAF signature from PDAC (hypergeometric test p-value = 0.066), IL1R and IL6 signaling have been implicated as key pathways used by iCAFs to promote an immunosuppressive tumor microenvironment.^51–53^

#### Vascular endothelial cells

As angiogenesis is a key contributor to the pathogenesis of PNETs, we subclustered 2,337 vascular endothelial cells from 14 non-functional, primary PNETs.^54,55^ cNMF resulted in 3 GEPs (**Fig. 3d-e**, **S6d-f**; **Methods**). We excluded one of the GEPs given lower feature number and higher percent mitochondrial genes, suggesting it consisted of lower quality nuclei (**Fig. S6e**). Endo-1 was enriched for terms related to glycosylation (*ADAMTS1/9*), and TGF-beta production and signaling (*SMAD3, HIF1A, TGFBR3, SULF1*). The expression of TGF-beta related genes and other genes related to mesenchymal features in Endo-1 (*FN1, CCN2, FBN1)* suggests a phenotype resembling endothelial to mesenchymal transition, a phenomenon of endothelial cell plasticity that is related to disease progression in multiple cancers.^56,57^ Endo-2 was enriched for terms related to VEGF-signaling, angiogenesis, and endothelial cell proliferation and showed increased expression of VEGF receptors (**Fig. 3e, S6f**).

#### Macrophages

Macrophages are the most abundant immune cell in the non-functional, primary PNET cohort and have been associated with more aggressive disease.^17–21^ We recovered 706 single-nucleus profiles and used cNMF to derive three GEPs (**Fig. 3f-g**, **S6g**; **Methods**). We compared the GEPs to previously described macrophage subsets derived from pan-cancer single-cell RNA-seq studies (**Fig. 3h**).^58,59^ The majority of TAMs belonged to the GEP Macro-1, which had significant overlap with the M2 signature (hypergeometric FDR = 0.006) and included canonical marker genes of M2 macrophages (*CD163, CLEC7A*). Although no GEPs had significant overlap with M1 macrophage or inflammatory TAM signatures, among macrophages with the highest activity of Macro-1, a subset also showed high expression of inflammatory markers (*TNF, IL1B, CCL3, CCL4*) (**Fig. S6h**). This is consistent with a published scRNA-seq study of gastroenteropancreatic NETs, which showed that macrophages in the GEP-NET TME may coexpress M1 and M2 markers.^21^ Two GEPs, combined to form Macro-2, had significant overlap with the lipid-processing TAM signature (hypergeometric FDR = 5.79×10^−3^ and 2.80×10^−5^, respectively), which included genes associated with immunosuppression and disease progression (*GPNMB*, *APOE*).^58,60^ Macro-3 had significant overlap with the angiogenic TAM signature (hypergeometric FDR = 2.80×10^−5^), a subset of TAMs that produce angiogenic growth factors in hypoxic tumor regions and are associated with worse prognosis and lack of response to immunotherapy in other cancer types.^55^ Overall, macrophages in the PNET TME appear to adopt a more immunosuppressive phenotype, with subsets showing elevated angiogenic and lipid-metabolizing signatures.

### Macrophage-derived glutamate supports an immunosuppressive environment and more aggressive behavior in cancer cells

To investigate how malignant cell state may impact interactions in the TME, we inferred cell-cell interactions between malignant cells expressing different GEPs and other cells in the TME using CellChat (**Fig. S7a**; **Methods**).^61^ Among the malignant GEPs, the two with the highest strength of predicted interactions with other cells were Malig-3 and Malig-4 (**Fig. S7a**). The interactions of Malig-3 cells were largely with other malignant subtypes via neurexin signaling (**Fig. S7a-b**), which are cell-adhesion molecules that mediate synapse formation and neuronal communication in the nervous system and insulin secretion from beta cells.^62,63^ These findings suggest that neurexins may also mediate interactions and adhesion between malignant cells in PNET.

As macrophage infiltration has been associated with more aggressive clinical outcomes in PNET,^17–19^ we next investigated how macrophages interacted with malignant cells. We found that macrophage-derived glutamate was predicted to signal to malignant cells via AMPA glutamate receptors, with highest probability of interaction with Malig-4 cells (**Fig. 4a, S7c**). Tumor specimens with a high proportion of Malig-4 cells showed upregulation of glutaminase and a glutamate transporter (*SLC1A3*) in macrophages, suggesting augmented glutamate production and export (**Fig. 4b**). This led us to ask if increased glutamate production may alter metabolism in macrophages and subsequent functional polarization, as previous studies have highlighted the importance of the citric acid cycle in supporting M2 macrophage polarization, an immunosuppressive subtype of macrophages.^64,65^ To this end, we used Compass to infer the activity of metabolic pathways in individual macrophages,^66^ and found that macrophages from samples with a high proportion of Malig-4 cells had significantly higher flux through glutamate metabolism reactions, such as glutamate dehydrogenase or alanine transaminase, which converts glutamate to alpha ketoglutarate, as well as reactions in the citric acid cycle (**Fig. 4c**). Furthermore, macrophages from these samples also had higher expression of M2 markers (**Fig. 4d**). We then asked if glutamate signaling may alter activity in malignant cells. Not only did we find that Malig-4 cells had the highest expression of AMPA glutamate receptors, but we discovered that these cells also had higher expression of downstream MAPK/ERK pathway members, which can be activated by AMPA receptor signaling in cancer cells (**Fig. 4e**), promoting the migratory and invasive capacity of cancer cells.^67,68^

**Figure 4.**
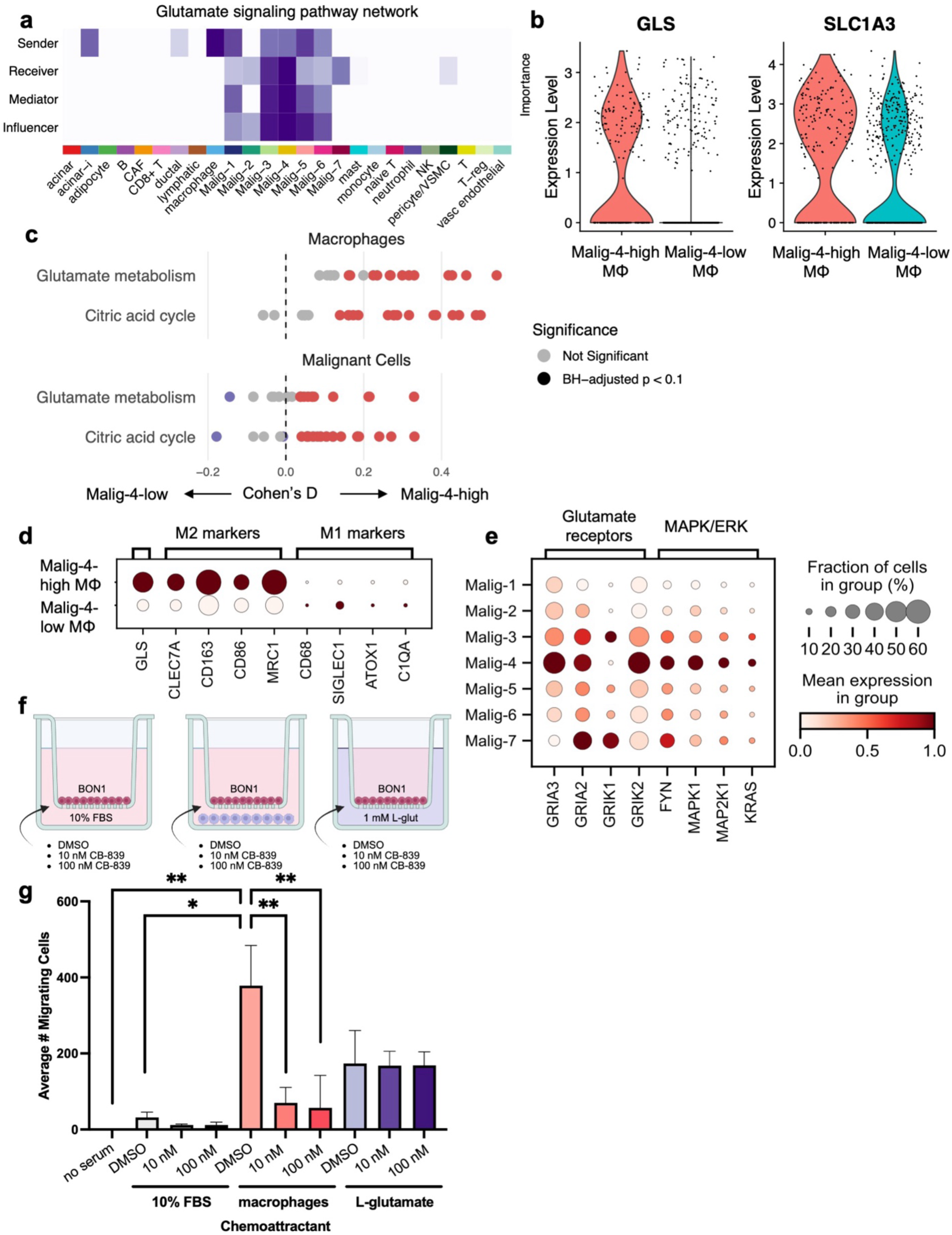
Glutamate metabolism supports immunosuppressive signaling in macrophages and invasive behavior in malignant cells. **(a)** Importance of different cell types (purple) in the signaling of glutamate in the tumor microenvironment, with macrophages as the most important senders (top row) and Malig-4 cells as the most important receivers (second row). Mediators (third row) are cell types with the highest betweenness centrality and influencers (fourth row) are the cell types with highest information centrality from CellChat network analysis. **(b)** GLS expression (left) and SLC1A4 (glutamate transporter, right) in macrophages from Malig-4-high samples (red) and Malig-4-low samples (teal). **(c)** Dotplot of Cohen’s D comparing metabolic pathway activity scores calculated by Compass for macrophages (top) and malignant cells (bottom). Red dots indicate the reaction has a significant Cohen’s d (BH-adjusted p < 0.1) and is upregulated in Malig-4-high cells; blue dots indicate the reaction has a significant Cohen’s D and is upregulated in Malig-4-low cells. **(d)** Dotplot of scaled average gene expression of M2 and M1 macrophage marker genes (columns) comparing macrophages from Malig-4-high and -low samples. **(e)** Dotplot of scaled average gene expression (columns) for malignant cells in each GEP (rows). **(f)** Schematic of migration assay of BON1 cells in response to various chemoattractants was determined using a transwell migration chamber (**Methods**). In brief, 100,000 serum-starved BON1 cells (red) were seeded in the upper chambers of the transwell inserts while the bottom chamber contained various chemoattractants: (1) 10% FBS; (2) 10% FBS with mouse-derived peritoneal macrophages (blue); or (3) 10% FBS with 1 mM L-glutamate. Bottom chambers were treated with 0.1% DMSO, 10 nM CB-839 (glutaminase inhibitor), or 100 nM CB-839. After 48 hours, migrating cells on the bottom surface of the insert were counted and averaged across 5 representative fields. **(g)** Results of migration assay. Y-axis represents average number of cells across 5 fields of view for each replicate. Two technical replicates and 3-5 biological replicates were performed for the no serum, FBS, and macrophage conditions and 1 technical replicate and 2 biological replicates were performed for the glutamate conditions. Error bars represent standard deviation, and significance was derived from pairwise Mann-Whitney U test. \**p* = 0.0357, \*\**p =* 0.0079.

To functionally validate this glutamate-mediated macrophage-cancer cell crosstalk inferred from our snRNA-seq data, we performed a transwell assay to assess the migratory potential of cancer cells in response to macrophages and glutamate (**Fig. 4f**). BON1 human PNET cells were plated on the insert and the chemoattractant was either mouse peritoneal macrophages or 10% FBS alone. We found that macrophages had a significantly stronger chemoattractant effect on BON1 cells compared to 10% FBS alone, but this effect was lost when the macrophages were treated with a glutaminase inhibitor (CB-839) (**Fig. 4g, S7d**), suggesting that the macrophage-mediated increase in BON1 cell migration was at least in part through glutamate production. When 1 mM of L-glutamate was used as chemoattractant, there was an intermediate amount of migration, while the addition of CB-839 did not alter this level of migration (**Fig. 4g, S7d**). This result is consistent with the mechanism of CB-839 in blocking the conversion of glutamine to glutamate without affecting extant glutamate. Overall, these findings support a potential mechanism in which macrophage glutamate production promotes aggressive behavior in PNETs by facilitating cancer cell invasion and driving M2 polarization to maintain an immunosuppressive microenvironment.

### Analysis of asynchronous primary PNET and liver metastases reveal limited genomic changes and increased VEGF-signaling throughout progression

Studying primary PNET tumors in isolation is limited by the lack of temporal dynamics. Thus, we were interested in profiling the transcriptomic and genomic landscape of an unusual case study from our cohort (P18). This patient had an untreated, primary PNET resected in 2015 (pT3N0 / grade 2), followed by a first metastatic event to the liver that was surgically resected in 2018, followed by a second metastatic event to the liver in 2021 that was also resected (**Fig. 5a**). This rare instance of three asynchronous and surgically resected primary and metastatic lesions over five years without any intervening neoadjuvant or adjuvant treatment afforded the unique opportunity to track transcriptional and genomic changes throughout the natural disease progression in this patient. To this end, we performed snRNA-seq and whole exome sequencing on frozen surgical resection specimens of the primary tumor (PT1), first metastatic lesion (MT1), second metastatic lesion (MT2), as well as whole exome sequencing of the patient’s peripheral blood leukocytes as a germline control.

**Figure 5.**
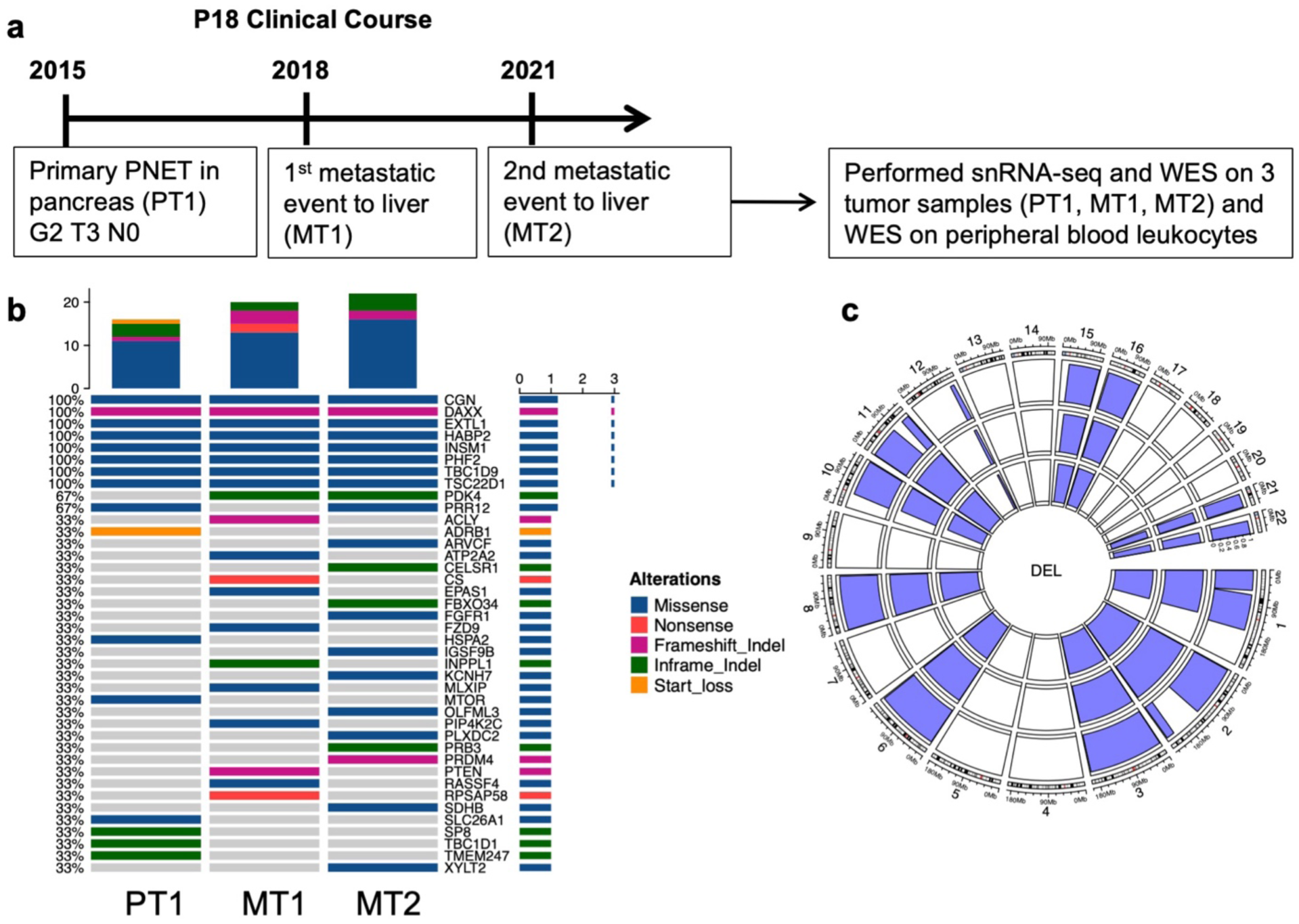
Case study of a primary nonfunctional PNET with two asynchronous metastases reveals a common clonal origin. **(a)** Clinical course of P18. **(b).** Oncoplot of somatic non-silent mutations (legend) of known oncogenes, tumor suppressors genes, and mutations predicted to be deleterious in two of three algorithms (SIFT, Polyphen2 HVAR, Polyphen2 HDIV) across samples (columns). Horizontal barplot represents the number of samples each mutation is present in and vertical barplot represents the number of mutations per sample shown on the oncoplot. **(c)** Somatic copy number alterations in PT1 (outer circle), MT1 (middle circle), and MT2 (inner circle) determined by FACETS on whole exome sequencing data with deletions shown in blue.

The genomic alterations across the three tumors were remarkably similar. The tumor mutational burden was overall low, consistent with what has been reported in PNETs,^7^ but increased throughout progression, from 0.813 coding mutations/Mb in PT1 to 1.122 coding mutations/Mb in MT1 to 1.432 coding mutations/Mb in MT2. The genetic landscape of somatic mutations was also remarkably similar throughout progression (**Fig. 5b**). Some notable mutations that were maintained in the primary and metastatic lesions include a missense mutation in *INSM1* (p.G258E) and frameshift deletions in *DAXX* causing early stop codons (p.L23Vfs*13, p.L98Vfs*13, p.L110Vfs*13). Inactivating mutations in *DAXX* are associated with ALT and more aggressive disease in PNET, consistent with this patient’s clinical course and the observation that the two metastatic lesions were positive for ALT (**Fig. 1a**).^14^ The only mutation gained in the metastatic samples compared to the primary tumor was an inframe insertion in *PDK4* (F43_G44insH), a gene involved in regulating pyruvate oxidation.^69^ The copy number alterations identified from both the whole exome sequencing data and inferred from the single-nucleus profiles were concordant and revealed that the copy number alterations were largely similar across tumors (**Fig. 5c S8a**). The copy number alteration profile was characterized by loss of heterozygosity in large parts of chromosomes 1, 2, 3, 6, 8, 10, 11, 15, 16, 21, and 22. This pattern of aneuploidy along with a mutation in *DAXX*, a gene involved in chromatin remodeling, is consistent with the Group 1 genomic subgroup associated with poor prognosis.^9^ Together, this analysis suggests that the primary tumor and two asynchronous metastases derive from the same genetic clone. Furthermore, it suggests that somatic mutations are not key drivers of metastatic progression in this patient.

Given the relatively stable mutational landscape, we next investigated whether the transcriptional landscape of the tumors changed throughout progression. After quality control, we recovered 16,836 single-nucleus profiles, consisting of 13,791 malignant and 3,045 nonmalignant cells (**Fig. 6a**). We annotated the malignant and non-malignant cell types as described previously (**Fig. S8b**; **Methods**). Of note, PT1 had a much lower cell count compared to the metastatic tumors and very few nonmalignant cells were identified. While this may represent the composition of the tumor microenvironment, this may also be because the sample was already six years old when sequencing was performed. Malignant cells formed the largest proportion of cell types in each sample (**Fig. S8c**). In PT1, epithelial cells formed the largest proportion of nonmalignant cell types, while in the metastatic tumors, the endothelial compartment formed the largest proportion of nonmalignant cells and increased proportionally from MT1 to MT2 (**Fig. S8b**). Macrophages were the most abundant immune cell type, and their proportion increased from MT1 to MT2.

**Figure 6.**
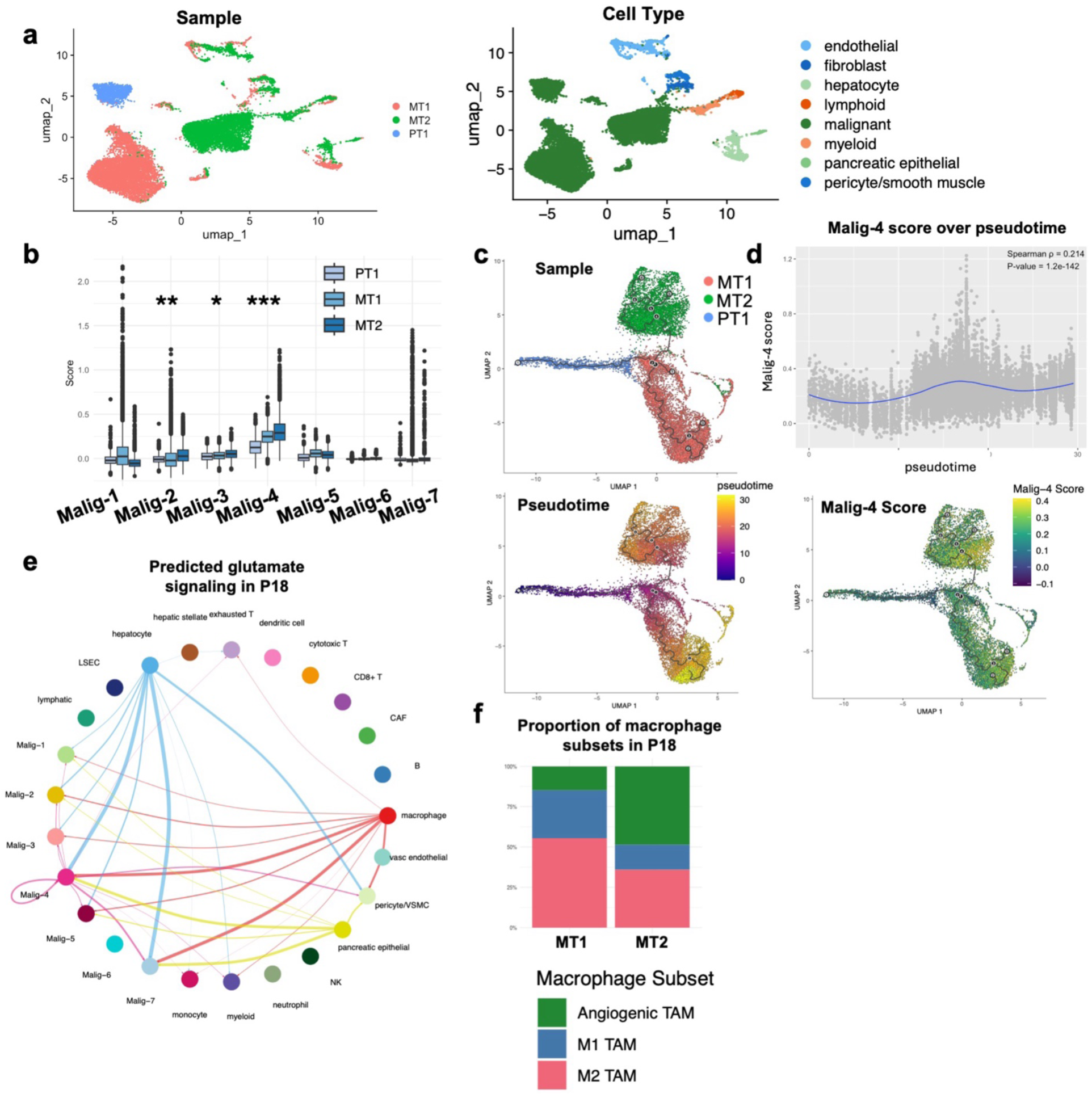
Transcriptomic landscape of a case study of a primary nonfunctional PNET with two asynchronous metastases reveals that the Malig-4 GEP activity increases throughout progression. **(a)** UMAP embedding of single-nucleus profiles of cells in P18 colored by sample (left) and cell type (right). **(b)** Boxplots of the score of each malignant GEP in the single nucleus profiles of the malignant cells in P18 colored by sample (legend). Asterisks indicate programs with FDR < 0.05 and predicted slope > 0.01. *** p < 0.001 and slope > 0.05, ** p < 0.001 and slope > 0.02, * p < 0.001 and slope > 0.01. **(c)** UMAP embedding of malignant cells in P18 colored by sample (top) and pseudotime (bottom). **(d)** Correlation of Cytoskeletal/VEGF-signaling GEP score in malignant cells of P18 over pseudotime (top) and plot of Malig-4 score on the UMAP (bottom). **(e)** Circle plot of predicted interactions in the tumor microenvironment of P18 from CellChat showing macrophages as primary senders of glutamate to malignant cells with thickness of edge representing the strength of the interaction. **(f)** Proportion of macrophage subtypes in P18 by sample.

Next, we scored the malignant cells from these samples for the seven malignant GEPs identified previously (**Fig. 6b**). A small cluster containing malignant cells from MT1 and MT2 scored highly for Malig-7, representative an actively proliferating subpopulation (**Fig. S8d**). Notably, in all three tumors, Malig-4, which features elevated VEGF-signaling, was the most highly scored malignant cell state. Furthermore, Malig-4 expression significantly increased throughout progression (p-value < 2.2×10^−16^, Jonckheere-Terpstra test) (**Fig. 6b**). To further investigate the transcriptional changes throughout progression at the single-cell level, we used trajectory pseudotime analysis with Monocle3.^70^ As expected from the genetic analysis, we identified a branching point from PT1 to MT1 and MT2, suggesting that both metastatic lesions derived from the primary tumor (**Fig. 6c**). Furthermore, the Malig-4 score increased across the trajectory (Spearman rho = 0.214, p-value = 1.2 × 10^−142^) (**Fig. 6d**). Together, these results suggest that the Malig-4 cell state may play a role in PNET progression and metastasis.

Given our previous findings of glutamate driving M2 polarization of macrophages and cancer cell invasion, we examined the phenotype of macrophages in P18. As no macrophages were observed in PT1, we excluded it from this analysis. First, we inferred cell-cell interactions in P18 and found that like the cohort of primary tumors, macrophages were predicted to be signaling to Malig-4 expressing cancer cells with glutamate (**Fig. 6e**). We scored macrophages from P18 with signatures derived from a pan-cancer single-cell atlas of myeloid cells and assigned macrophages to the signature with the highest score.^59^ We found that while M2 macrophages were most abundant in MT1, the proportion of angiogenic TAMs, a subtype of macrophages associated with worse prognosis, is significantly increased in MT2 (Chi-square test of proportions p-value = 1.07×10^−6^) (**Fig. 6f**). Thus, these results suggest that the interactions between macrophages and the Malig-4 malignant cell state may play a key role in promoting more aggressive behavior in nonfunctional PNETs.

## Discussion

In this study, we use snRNA-seq to profile the largest cohort of PNETs to the best of our knowledge. While previous studies have used scRNA-seq to profile PNETs, we were able to recover a much higher percentage of malignant neuroendocrine cells compared to previous studies, either due to differences in sample preparation or differences in the cellular composition of the tumors in our cohort (**Fig. 1d**).^21,25^ With this larger number of malignant cells, we developed a *de novo* classification of malignant cell states based on GEPs. Our classification included Malig-5, which recapitulates a previously described beta cell-like state that is correlated with better overall survival in an independent bulk RNA-seq cohort.^11,12,15^ We also discovered two novel cell states: Malig-3, which is enriched for neuronal/synaptic signaling genes, and Malig-4, which was enriched for VEGF signaling and cytoskeletal genes. In our primary cohort, Malig-3 was significantly enriched in PNETs with more aggressive disease, ALT-positivity, and broad loss of heterozygosity in ten chromosomes, a genomic group associated with worse prognosis. Furthermore, high Malig-3 score in independent bulk transcriptomic datasets was significantly associated with worse overall survival. Malig-3 overlaps with the Metastatic Like Precursor bulk transcriptomic signature (MLP) previously associated with worse prognosis in PNETs,^8^ including shared expression of glutamate receptors (AMPA and kainate receptors), synaptic scaffolding genes (neurexins), ion channel and transport genes (*KCN* and *SLC* family genes), as well as genes related to stemness and neuronal development (*DCLK* family genes, *SOX11, SOX4*). Our cell-cell interaction analyses suggest that malignant cells expressing the Malig-3 GEP largely interact with other malignant cells via neurexins, a cell adhesion molecule commonly found in the nervous systems and pancreatic beta cells. Pseudo-synapses between cancer cells and neurons have been identified in multiple cancer types, such as breast cancer,^71^ and malignant neuroendocrine cells in small cell lung cancer and PNET model systems have been shown to have intrinsic electrical activity.^72,73^ Future studies should investigate potential neuronal mimicry and the formation of pseudo-synapses or other junctional interactions in PNET as a form of cell-cell communication that may promote tumorigenesis and progression in PNET.

VEGF signaling, which is enriched in the Malig-4 GEP, is an important pathway in PNET as several multikinase inhibitors that target VEGFR, such as the recently FDA-approved drug cabozantinib, improve progression free survival in patients with advanced disease.^55,74^ However, resistance to these drugs is widespread, with common mechanisms including tumor hypoxia via HIF1A activation or VHL inactivation, and alternative vascularization, such as through FGF or angiopoietin/Tie2 signaling.^75–77^ Our findings suggest that immune interactions in the microenvironment may play a role in disease progression in patients who have a large proportion of Malig-4 cells. Malig-4 rich specimens featured macrophages with higher expression of immunosuppressive markers, and angiogenic macrophages were more prevalent in the second metastatic lesion compared to the first metastatic lesion in our case study of P18. Furthermore, our predictions from patient-derived snRNA-seq data and *in vitro* results suggest a circuit by which preferential signaling interactions between malignant cells and macrophages both promote an immunosuppressive state in macrophages and greater migratory behavior in malignant cells. Suppressing macrophage-derived glutamate reduced the migration of PNET cancer cells, although glutamate in the TME may also derive from other cell types. Indeed, previous research has found that malignant PNET cells can hijack glutamate signaling pathways to promote proliferation and invasion via NMDA receptors.^78,79^ Our results suggest that targeting glutamate metabolism may be efficacious in PNET by inhibiting both cancer cell-intrinsic and -extrinsic protumorigenic effects. Furthermore, our results suggest that profiling malignant cell composition could improve personalized treatment in PNET. For example, a patient with high levels of the VEGF-signaling cell state may be an ideal candidate for a multi-kinase inhibitor, and future studies should explore combination treatment with a therapeutic that targets the immunosuppressive or angiogenic features of macrophages to improve efficacy and combat treatment resistance.

Finally, we presented a case study of an untreated non-functional primary PNET and two untreated asynchronous hepatic metastases over the span of six years. This rare exemplar provides a unique opportunity to track genomic and transcriptomic changes throughout disease progression. This patient had several genomic features suggestive of poor prognosis, including frameshift deletions in *DAXX,* ALT-positivity, and broad loss of heterozygosity in chromosomes 1, 2, 3, 6, 8, 10, 11, 16, 21, 22. Notably, there were very few somatic single nucleotide variants or copy number changes among the three tumors, strongly suggesting a monoclonal origin. These results are consistent with previously reported genomic profiling of paired primary and metastatic PNETs in which genomic changes are limited, suggesting that genomic alterations are not the main drivers of progression and metastasis in PNETs.^9^ The lack of clear genetic drivers in the metastatic tumors suggests an important role for non-coding mutations and/or non-genetic events. Possible causes highlighted by our transcriptomic analysis include expression of Malig-4 increased throughout progression, as well as an increased proportion of macrophages polarized towards the M2 and angiogenic phenotypes. Although Malig-4 was not found to be prognostic in the larger independent bulk-RNA-seq cohorts, this case study illustrates that this GEP may still contribute to disseminated progression, potentially through modulation of malignant-immune interactions in the TME.

Our study has several limitations. Although this is the largest cohort of human PNET analyzed by single-cell/nucleus transcriptomics to the best of our knowledge, the overall sample size was still relatively low. While we did find that Malig-3 was enriched in PNETs with aggressive disease in our primary cohort and in PNETs with lower overall survival in independent bulk RNA-seq cohorts, future studies should apply these signatures to larger sc/snRNA-seq datasets. Our survival analyses of the snRNA-seq derived GEPs in bulk transcriptomic cohorts may be confounded by expression of GEP genes in nonmalignant cells. Additionally, while previous groups have determined genomic categories of PNET based on both patterns of aneuploidy and somatic mutations of various genes (*ATRX, DAXX, MEN1*), since we did not have genetic data from patients except P18, we solely categorized the tumors based on patterns of aneuploidy determined by inferred copy number variants. Another limitation is that we were only able to profile two PNETs with neoadjuvant treatment. While one patient had higher levels of the beta cell-like GEP (Malig-5), corresponding to a better clinical course, the other patient did not show enrichment of any of the GEPs identified from the untreated subset of our cohort. This highlights the need to profile larger numbers of PNETs with treatment to understand how treatment affects the landscape of malignant cell states in PNET. Finally, in this study we were able to dissect the heterogeneity within and signaling among cell types in the TME of PNET with snRNA-seq. However, an inherent limitation of this approach is the cells are dissociated and there is a lack of spatial context. Future studies should take advantage of recent advances in spatial multi-omics to gain a greater understanding of cellular communities and signaling interactions with the added context of *in situ* relationships.

In summary, this study presents the largest transcriptomic dataset of human PNETs at single-cell resolution to date. Our refined taxonomy of malignant cell states identified two novel malignant GEPs: one enriched in neural/synaptic signaling genes that is associated with worse survival in independent cohorts, and one enriched for VEGF-signaling that was associated with progression in a case study of a metastatic PNET. We also identified signaling interactions in the TME between TAMs and malignant cells via glutamate, suggesting mechanisms by which an immunosuppressive environment may be maintained, and malignant cells can augment their invasive behavior. Together, this study provides an important resource with new insights into the diversity and biological relevance of malignant cell states in PNET and their interactions with the microenvironment.

## Methods

### Ethics

All patients in this study were consented without compensation to DF/HCC 02-240 and/or excess tissue biobank protocol 2022P001264, which were reviewed by the Massachusetts General Hospital (MGH) Institutional Review Board. This study is compliant with all relevant ethical regulations.

### Human patient specimens

For inclusion in this study, patients were required to have a diagnosis of pancreatic neuroendocrine tumor with a plan for surgical resection with or without prior neoadjuvant treatment (**Table S1**). Resected tumor samples were examined by a board-certified pathologist to confirm the diagnosis of PNET and then snap frozen and stored at −80°C for up to 6 years prior to processing.

### Assessment of Alternative Lengthening of Telomeres (ALT)

Representative slides from formalin-fixed paraffin-embedded blocks (FFPE) were reviewed and examined by a by a board-certified pathologist to confirm the presence and optimal view of PNET. Next, 5 µm FFPE sections were cut from the selected block to be used for ALT staining. ALT was assessed using a telomere-specific fluorescence in situ hybridization (FISH) assay on whole tumor sections, as previously described.^80,81^ Deparaffinized slides were hydrated through a graded ethanol series following paraffin removal with xylene. Antigen retrieval was performed with citrate buffer (pH 6.0; Cat# H-3300, Vector Laboratories). Slides were then hybridized overnight with a Cy3-labeled peptide nucleic acid (PNA) probe complementary to telomeric DNA repeats (Cat# F1002, Panagene). To verify hybridization efficiency, an Alexa Fluor 488-labeled PNA probe targeting centromeric DNA repeats (Cat# F3012, Panagene) was included in the hybridization solution. Nuclei were counterstained with DAPI (4′,6-diamidino-2-phenylindole; Sigma-Aldrich) and mounted with ProLong antifade mounting medium (Invitrogen). Cases were classified as ALT-positive based on previously established criteria^82^ which included the presence of large, ultrabright intranuclear telomeric foci in ≥ 1% of tumor cells and individual foci exhibiting signal intensities at least 10-fold greater than telomeric signals observed in non-neoplastic cells. Of note, areas of necrosis were excluded from evaluation.

### Nucleus isolation from frozen samples and single-nucleus RNA-seq

Nuclei isolation was performed as described previously using a protocol specific for pancreatic tumors.^22^ Approximately 8,000-10,000 nuclei per sample were loaded into each channel of a Chromium single-cell 3’ chip (V3, 10x Genomics) according to the manufacturer’s instructions. Single nuclei were partitioned into droplets with gel beads in the Chromium Controller to form emulsions, after which nuclear lysis, barcoded reverse transcription of mRNA, cDNA amplification, enzymatic fragmentation, and 5’ adaptor and sample index attachment were performed according to manufacturer’s instructions. The snRNA-seq sample libraries were sequenced on the NovaSeq 6000 (Illumina) with the following paired end read configuration: read 1, 28 nt; read 2, 90 nt; index read, 10 nt.

### snRNA-seq data pre-processing

BCL files were converted to FASTQ using bcl2fastq2-v2.20. CellRanger v6.0.0 with default parameters was used to demultiplex the FASTQ reads, align them to the hg38 human transcriptome (pre-mRNA) reference and extract the UMI and nuclei barcodes. The output of this pipeline is a digital gene expression (DGE) matrix for each sample, which has quantified for each nucleus barcode the number of UMIs that are aligned to each gene. For each sample, doublets were identified and removed using scDblFinder v.1.22.0 with default parameters and nuclei were filtered for minimum number of features of 200, proportion of mitochondrial genes and maximum number of features with thresholds determined for each sample. UMI counts were normalized by the total number of UMIs per nucleus and converted to transcripts-per-10,000 (TP10K) as the final expression unit.

### Dimensionality reduction, clustering and annotation

The treatment-naive primary non-functional tumors (n = 15) were aggregated into a single dataset; the tumors with neoadjuvant treatment (n = 2) were aggregated into a single dataset; and the paired primary and two metastatic samples from P18 (n = 3) were aggregated into a single dataset. The log2(TP10K + 1) expression matrix was constructed for downstream analyses. We identified the top 2,000 highly variable genes with Seurat v.5.2.1 FindVariableFeatures function and scaled the data with the ScaleData function. We performed principal component analysis over the variable features and identified the top 50 principal components. We built a k-nearest neighbors graph based on the top 50 principal components and performed community detection on the neighborhood graph with the Louvain graph clustering method using the functions FindNeighbors and FindClusters. Individual nucleus profiles were visualized using Uniform Manifold Approximation and Projection (UMAP) which was calculated with the function RunUMAP. To annotate cells, first we identified malignant and nonmalignant compartments (i.e. stromal, immune, epithelial) by examining the expression of known cell type-specific gene expression signatures. For nonmalignant cell annotation, each nonmalignant cellular compartment was subsetted and variable genes, scaling, principal components, UMAP, and Louvain clusters were recalculated. Batch correction based on sample ID was performed on the selected number of principal components with Harmony v1.2.3. Then distinct populations of nonmalignant cells were determined within each subcluster by the expression of cell type-specific marker genes. During the annotation process, clusters of nuclei showing strong co-expression of cell type-specific marker genes and neuroendocrine-specific genes were determined to be doublets and manually removed.

### Inferring copy number variations from single nucleus profiles

The malignant cells were confirmed by running inferCNV v1.3.6 jointly on all malignant and nonmalignant single nuclei for each patient to confirm that the designated malignant cells had a higher copy number burden compared to the nonmalignant cells.^83^ Default parameters were used, including a 100-gene window in sub-clustering mode and a six-level hidden Markov model to predict the copy number variant in each nucleus. Next, each tumor was classified into previously defined genomic groups based on the deletions identified in inferCNV.^9^ A tumor was classified as Group 1 (loss of heterozygosity of chromosomes 1, 2, 3, 6, 8, 10, 11, 16, 21, 22) if it contained deletions in all ten chromosomes; a tumor was classified as Group 2 if it contained a deletion in chromosome 11, but not in all other ten chromosomes; and otherwise in Group 3 (variable aneuploidy).

### Consensus non-negative matrix factorization

We inferred gene expression programs (GEPs) among the malignant, fibroblast, vascular endothelial, and macrophage compartments in primary, non-functional untreated PNETs with non-negative matrix factorization implemented in the package cNMF v1.7.^84^ For the fibroblast, vascular endothelial, and macrophage compartments, only coding genes were used. With this method, the gene expression matrix is factorized into two matrices: one defining the genes that constitute a GEP and the other defining the activity of each GEP in a nucleus. We repeated NMF 200 times per cell type category and computed a set of consensus programs by aggregating results from all 200 runs and computed a stability and reconstruction error. To determine the optimal number of programs (k) for each cell type and condition, we chose the cNMF solution that maximized stability and minimized error, while maintaining biological coherence. Each GEP was defined by the top 100 weighted genes and annotated by a combination of performing EnrichR analyses with established gene sets and comparison to known signatures.^12,26,28,58,59^ To determine the similarity to known signatures, a two-sided hypergeometric test was performed to quantify overlap and p-values were adjusted with Benjamini-Hochberg procedure.

For the malignant compartment, we focused on GEPs that were expressed across multiple patients and that were robust to downsampling of the malignant cells. To perform the downsampling, we randomly selected 20%, 40%, 60%, 80%, and 90% of malignant cell barcodes and performed cNMF with the same number of programs in the optimal solution when using 100% of the malignant cells (k = 25). Then we determined the Jaccard similarity between the 100 weighted genes of the downsampled GEPs and GEPs determined with 100% of the malignant cells and retained 22 GEPs with greater than 70% Jaccard similarity at downsampling to 90% of malignant cells. We excluded 15 GEPs that were patient specific, resulting in 7 GEPs for downstream analysis. To assign malignant cells to a particular GEP, we performed L1 normalization across all 7 GEPs such that the vector of GEP activities for a nucleus summed to 1. The nucleus was then assigned to the GEP with the highest score.

### Inferring gene regulatory networks of malignant gene expression programs

The activity of gene regulons in each nucleus annotated as malignant was inferred using the pySCENIC v. 0.12.1 pipeline.^29^ The GRNBoost2 algorithm was used to infer co-expression modules based on transcription factor and target gene relationships. Then candidate regulons were identified by mapping co-expression modules to DNA motif enrichment using reference motif databases for the human hg38 SCENIC motif collection v10 and the corresponding HGNC Motif2TF v10 annotation. Finally, the activity of each regulon in individual nuclei was calculated using AUCell. Next, we calculated the Pearson correlation between the regulon activity and malignant GEP score to identify the most highly correlated regulons for each malignant GEP.

### Determining enrichment of malignant cell states in different genomic groups, clinical outcomes, and ALT status

Tumors were classified into genomic groups as described above and into clinical groups determined by the clinical behavior. Aggressive clinical behavior was defined as a G3 primary tumor, nodal metastases, or future metastasis (n = 5) and otherwise was defined as benign clinical behavior (n = 9). Tumors were classified into ALT-positive (n = 4) or negative status (n = 10) as described above. To evaluate the association between each genomic or clinical group and GEP scores, a Gamma mixed effects regression model with a log link was implemented with the glmmTMB v. 1.1-11 package in R. A Gamma mixed effects regression model was deemed appropriate as GEP scores derived from cNMF were positive and right-skewed. A negligible constant (e = 1E-6) was added to each GEP score so that all outcomes were > 0. For each GEP, a separate model was estimated using Laplace approximation with GEP score as the dependent variable; genomic, clinical group, or ALT status as a fixed effect; and sample ID as a random effect to account for the non-independence of cells deriving from the same sample. Wald *z*– statistics provided two-sided *p*-values for each fixed-effect contrast. To control for multiple comparisons, FDR adjustment was applied to p-values with the Benjamini-Hochberg procedure.

### Survival analysis with bulk RNA-seq data

Bulk RNA-seq data from three previously published PNET cohorts with overall survival annotated were obtained.^6,12,15^ Only primary resected PNETs were included in the analysis, yielding 140 patients for further analysis (PAEN-AU, n = 32; Chan *et al.,* n = 24; EGA, n = 84). Processed log2(transcripts per million) were available for the Chan and PAEN-AU datasets, while for the EGA dataset, BAM files were aligned to the hg19 genome and quantified with FeatureCounts v2.0.2 and then converted to log2(transcripts per million).

All survival analyses were performed using the survival v.3.8.3 package in R. Each malignant GEP was analyzed separately. For each tumor, the GEP score was determined by summing the expression of the top 100 weighted genes for each GEP and then performing z-score normalization within each cohort independently to account for batch effects. Tumors were stratified into two groups based on GEP score: scores higher than the median of the cohort were classified as “high” and scores below the median of the cohort were classified as “low”. Overall survival (days) was analyzed with the Kaplan Meier method. Statistical comparison between groups were performed using a log-rank test (rho = 1).

### Scoring gene signatures for each single-nucleus profile

To score the malignant cells from the PNETs with neoadjuvant treatment (P17 and P18) and the paired primary and two metastatic samples from P18 with the malignant GEPs, we used the AddModuleScore function in Seurat. We calculated the average expression level for the 100 genes that defined each GEP subtracted by the aggregated expression of a set of 100 control genes that were randomly selected from genes that had similar average expression. We used the same approach to calculate the score of M1, M2, and angiogenic tumor associated macrophage signatures in nuclei annotated as macrophages from P18.^59^ Macrophages were assigned to either M1, M2, or angiogenic macrophage based on the maximum score.

### Inferring activity of metabolic reactions in single nuclei

To infer the activity of metabolic reactions in each cell, we use Compass v.0.9.10.2 with python 3.8.13 and Cplex version 20.10. Compass infers the activity of metabolic reactions in Recon2 based on the log normalized expression of enzyme-coding genes.^66,85^ First, Compass returns a reaction penalty matrix where the penalty is inversely correlated with reaction likelihood (i.e. a high score means a low likelihood of a reaction happening). We converted the reaction penalties to reaction scores with the function get_reaction_consistencies. Then, we compared the reaction scores of malignant cells with high levels of Malig-4 (GEP score > 0.1) compared to low levels of Malig-4 (GEP score < 0.1). We also compared the reaction scores of macrophages from primary, non-functional, untreated samples with a high level of Malig-4 (defined as proportion of Malig-4 activity > 0.1, n = 5)) and a low level of Malig-4 activity (n = 9). Comparison between the two groups was determined by a two-sided Wilcoxon test with adjusted p-value correction with the Benjamini-Hochberg procedure.

### Inferring cell-cell interactions

We used CellChat v.2.1.0 and the human CellChat database to infer interactions between cell types and malignant cell states in our snRNA-seq dataset.^61^ CellChat models ligand-receptor interactions with the law of mass action based on the expression levels of ligands, receptors, and cofactors. We used the “triMean” method to determine statistically significant reactions, which is the most stringent method.

### Cell culture

The human PNET cell line, BON1, was derived from a pancreatic serotonin-secreting neuroendocrine tumor^86^ and was obtained from Christopher Heaphy. BON1 was cultured in DMEM supplemented with 10% fetal bovine serum (FBS) and 1% penicillin/streptomycin. Cells tested negative for mycoplasma contamination and cells under passage 25 were used.

### Isolation of peritoneal macrophages from wild-type mice

Wild-type C57BL/6J mice of at least 10 weeks of age were purchased from Jackson Laboratory and used to isolate peritoneal macrophages. All mice were housed in accordance with the institutional guidelines. All procedures involving mice were approved by the institutional animal care and use committee of Massachusetts General Hospital. Peritoneal macrophages were isolated from the mice as described previously.^87^ Briefly, mice were euthanized and ice cold DMEM supplemented with 10% FBS and 1% penicillin/streptomycin was injected into the peritoneal cavity and then collected and centrifuged to pellet the cells. The cell pellet was resuspended in ice-cold DMEM and 1 million cells were plated in each well of 24-well plates.

### Migration transwell assay

BON1 cells were serum-starved in DMEM for 24 hours. Peritoneal macrophages were isolated and allowed to settle onto a 24-well plate overnight. The next day, macrophages were washed thoroughly to remove any nonadherent cells and replaced with DMEM containing 10% FBS with DMSO, 10 µM, or 100 µM of CB-839 (Selleck, S7655). Then, chambers with 12 µm porous polycarbonate membranes (Millipore, PIXP01250) were applied to the 24-well plates and 100,000 BON1 wells were seeded in serum-free DMEM in each chamber. For conditions without macrophages, the wells were filled with serum-free DMEM with 0.1% DMSO, DMEM with 10% FBS with 0.1% DMSO, 10 nM, or 100 nM of CB-839, or DMEM with 10% FBS and 1 mM L-glutamate (Abcam, ab120049) with 0.1% DMSO, 10 nM, or 100 nM of CB-839. After 48 hours of incubation at 37°C, BON1 cells that migrated to the bottom chambers were fixed, stained with Differential Quik III stain kit (Electron Microscopy Science, 26096-25) according to manufacturer’s protocol and counted in five fields under 10X magnification (Nikon Ts2-FL). Cells were counted in QuPath^88^ using cell detection with default parameters and a minimum intensity of 0.5. For each replicate, the cell count across the five fields were averaged to determine the average number of migrating cells. Significance was determined by a Mann-Whitney U test.

### Whole-exome sequencing and processing

Whole-exome sequencing was performed on genomic DNA extracted from frozen tumor samples from P18 of a primary tumor and two paired, asynchronous liver metastases (PT1, MT1, and MT2), as well as genomic DNA extracted from frozen peripheral blood leukocytes from P18 that were isolated at the same time of the surgical resection of the MT1. The genomic DNA was sent to Novogene for whole exome sequencing with Novaseq 6000 (Illumina). Reads were mapped to the reference hg38 genome and output into bam format with the Burrows-Wheeler Aligner v0.7.17, Samtools v1.8. was used to sort the bam file, and Picard v2.18.9 was used to merge all bam files from the same sample and mark the duplicated reads. Mutect2 v4.1.8.1 workflow on Terra was used to identify germline versus somatic single nucleotide variants (SNV) and insertions/deletions (Indels) and the web implementation of ANNOVAR (wANNOVAR) was used to annotate the somatic mutations.^89,90^ Non-silent somatic SNV/Indels in known oncogenes and tumor suppressors or predicted to be deleterious in 2 of 3 algorithms (SIFT, Polyphen2 HVAR, Polyphen2 HDIV) were plotted on an oncoplot with ComplexHeatmap v2.20.0. Somatic copy number alterations were determined by converting the bam files to pileup files with snp-pileup v0.6.1, followed by using the R package FACETs v0.6.2.^91^

### Pseudotime trajectory analysis

To infer pseudotime from the single nucleus profiles of the tumor samples from P18, we used Monocle3 v.1.3.7 with default parameters.^70^ First, we subset only the malignant cells from PT1, MT1, and MT2. Then we used Monocle3’s implementation of dimensionality reduction with PCA and UMAP for visualization, followed by clustering with the Leiden algorithm. The trajectory graph was learned using reversed graph embedding and pseudotime was assigned by selecting the malignant cells from PT1 as the root and computing distances along the inferred trajectory. Spearman correlation was calculated between Malig-4 score (calculated by AddModuleScore as described above) and pseudotime.

## Data Availability

Cell Ranger unfiltered and filtered barcode counts are uploaded to GEO (GSE301075, reviewer token: qdexyaaodbsftiv). The code used for the bulk of the scRNA-seq analysis is available on Zenodo (https://doi.org/10.5281/zenodo.15770193). Publicly available bulk RNA data that was analyzed in this study are available on in the European Genome-Phenome Archive EGAD00001006991; the Pancreas Genome-Phenome Atlas [PAEN-AU](https://pancreasexpression.org/analytics/cohort/icgc_paen_au/); and the Gene Expression Omnibus GSE118014.

## Conflict of Interest

All authors declare no interests related to this work.

## Acknowledgements

We are grateful to the patients and families who contributed their time and specimens to this study. We would like to acknowledge Ella Perrault, George Eng, Jorge Raldon, and Jamie Barth for helpful discussions and technical assistance. This work was supported by the Neuroendocrine Tumor Research Foundation (W.L.H.), Ruane Family Foundation Award in Endocrine and Neuroendocrine Tumors (W.L.H., M.H.), NCI R21CA286402 (C.M.H.), and Department of Defense Rare Cancers Research Program HT9425-23-1-0819 (C.M.H.). The funders had no role in study design, data collection and analysis, decision to publish or preparation of the manuscript.

## Notes

### Competing Interest Statement

The authors have declared no competing interest.

