## Supplemental Figures for "Distinct malignant cell states and myeloid glutamate signaling associated with aggressive pancreatic neuroendocrine tumors"

### Supplementary Figures

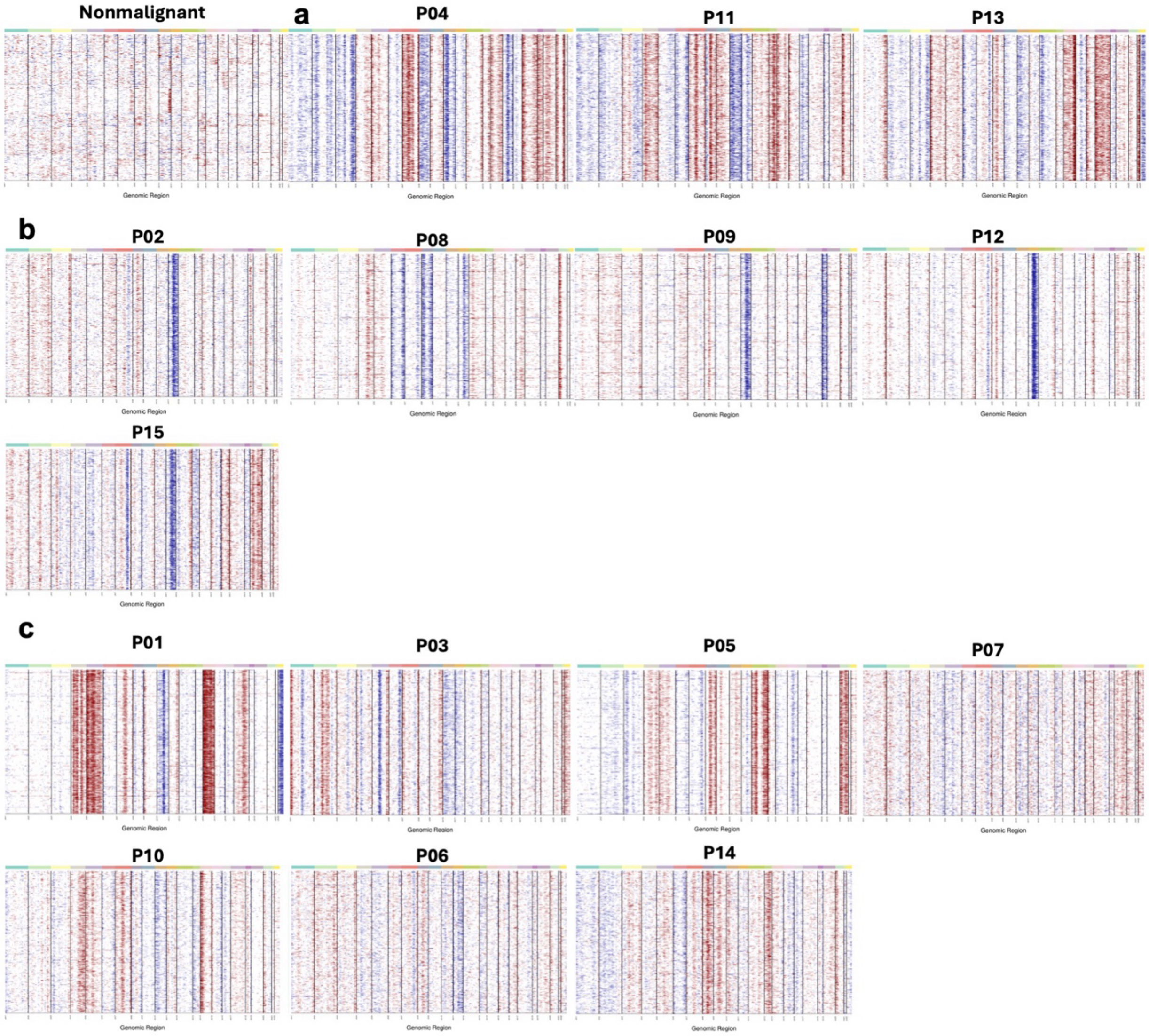


**Figure S1. Inferred copy number variations partition PNETs into different genomic subgroups.** Inferred amplifications (red) and deletions (blue) based on expression of a sliding 100-gene window in each chromosomal locus (columns) for each cell (rows) compared to the nonmalignant cells (labeled). **(a)** Group 1 tumors with broad loss of heterozygosity in chromosomes 1, 2, 3, 6, 8, 10, 11, 16, 21, 22. (**b)** Group 2 tumors with loss of chromosome 11. (**c)** Group 3 tumors with variable aneuploidy.

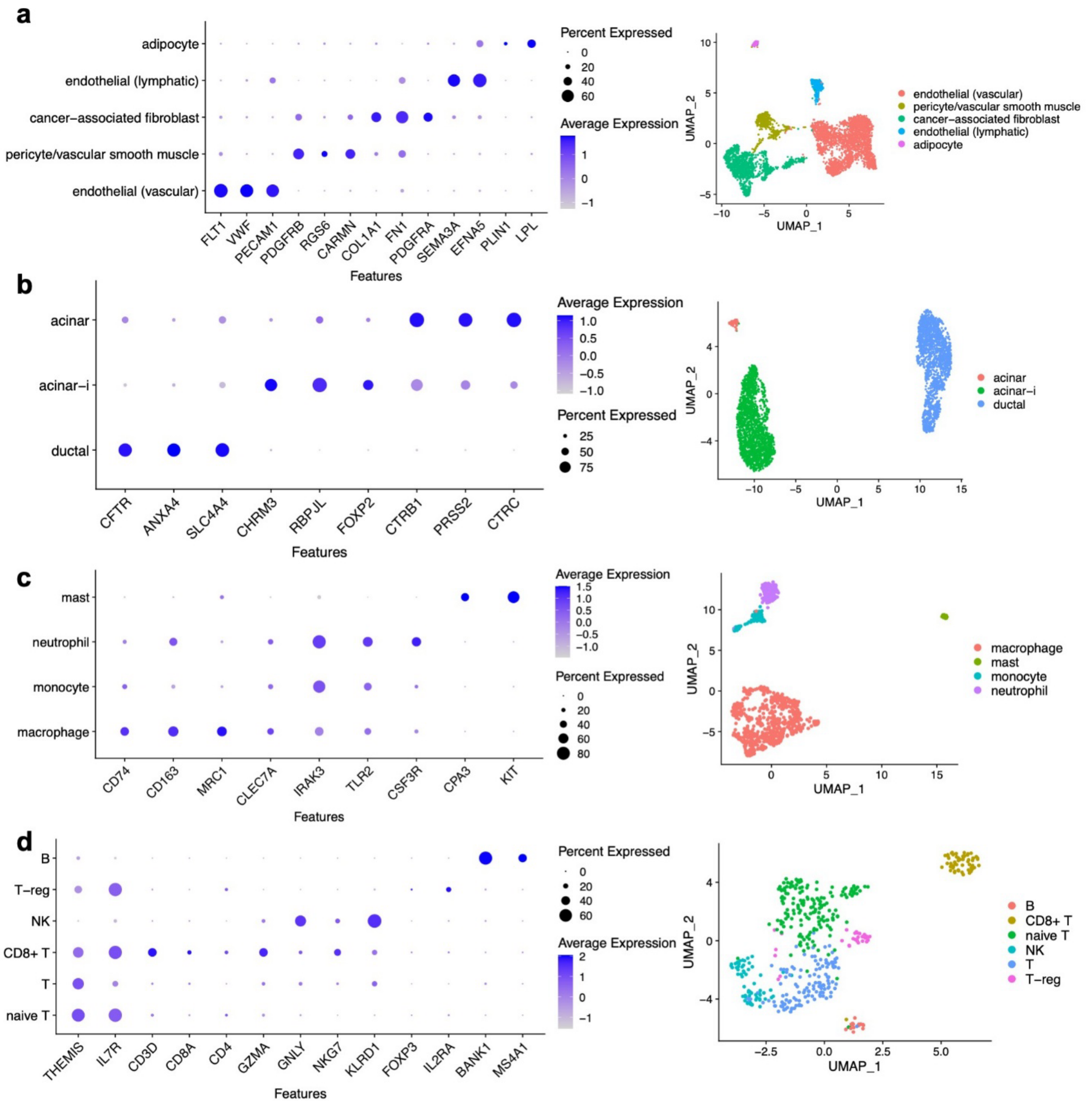


**Figure S2. Annotation of nonmalignant cell types in untreated primary PNET.** Average expression (color spectrum) and percent of cells expressing (dot size) selected marker genes (columns) across annotated cell types (rows) in **(a)** the stromal compartment; **(b)** the nonmalignant epithelial compartment; **(c)** the myeloid compartment; and **(d)** the lymphoid compartment. The size of each dot represents the percent of cells in the cell type expressing the gene.

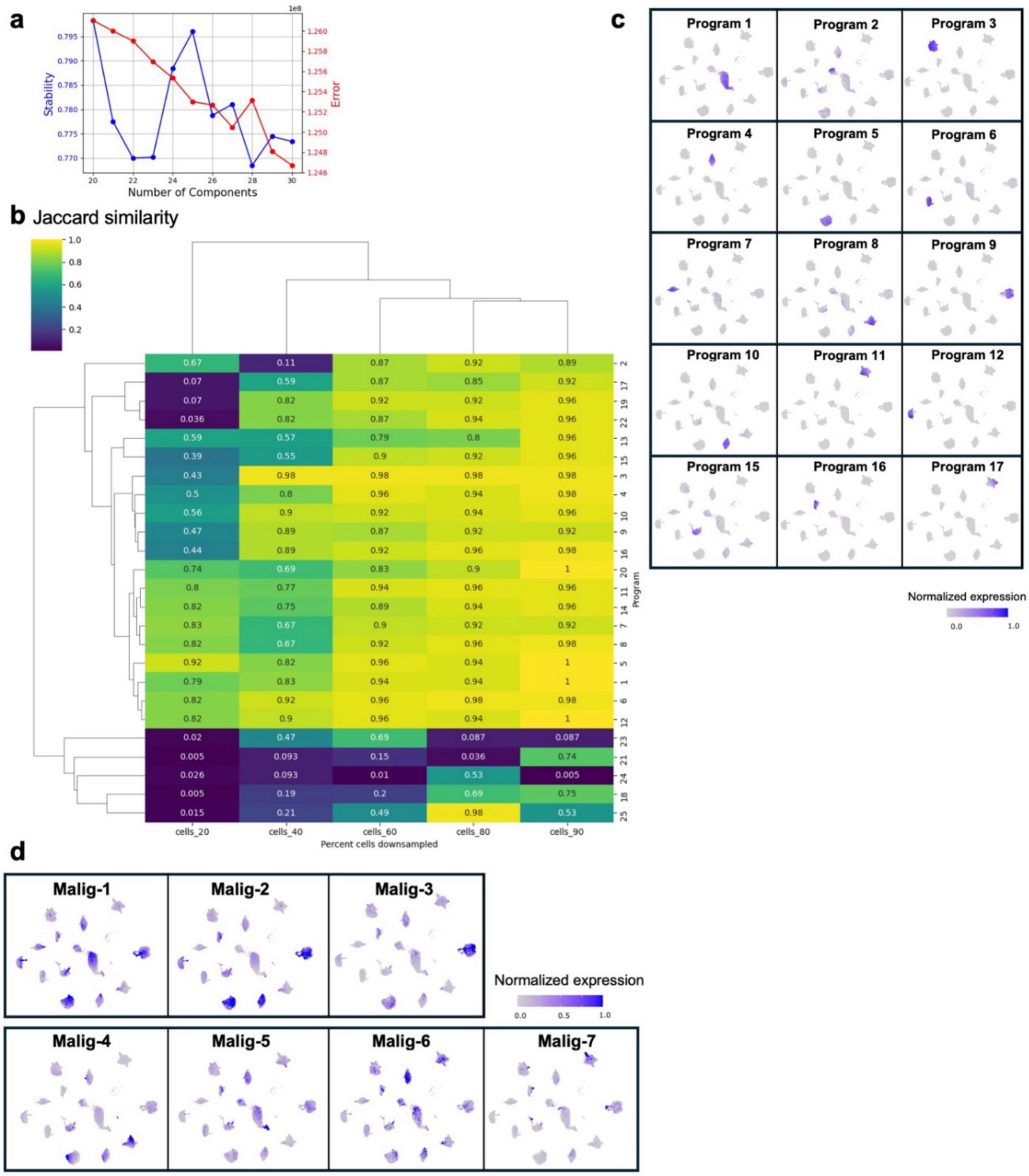


**Figure S3. Consensus NMF identifies GEPs shared across patients with nonfunctional, primary PNET. (a)** Estimated stability (blue, left y-axis) and error (red, right y-axis) in the cNMF solution (left) learned with different programs (components, x-axis) for malignant cells. **(b)** UMAP embedding of single nucleus profiles (dots) of malignant cells from primary, nonfunctional, untreated PNET tumors colored by the normalized expression score (blue) of each patient-specific program. **(c)** Heatmap showing Jaccard similarity (legend) between programs identified with downsampling of 20%, 40%, 60%, 80%, and 90% of malignant cells (columns) with the original 25 programs identified by cNMF of all malignant cells. 23 of the 25 programs were robust to downsampling of 90% of cells. **(d)** UMAP embedding of single nucleus profiles (dots) of malignant cells from primary, nonfunctional, untreated PNET tumors colored by the normalized expression score (blue) of each patient-distributed program. Color scale saturates at the 99th percentile.

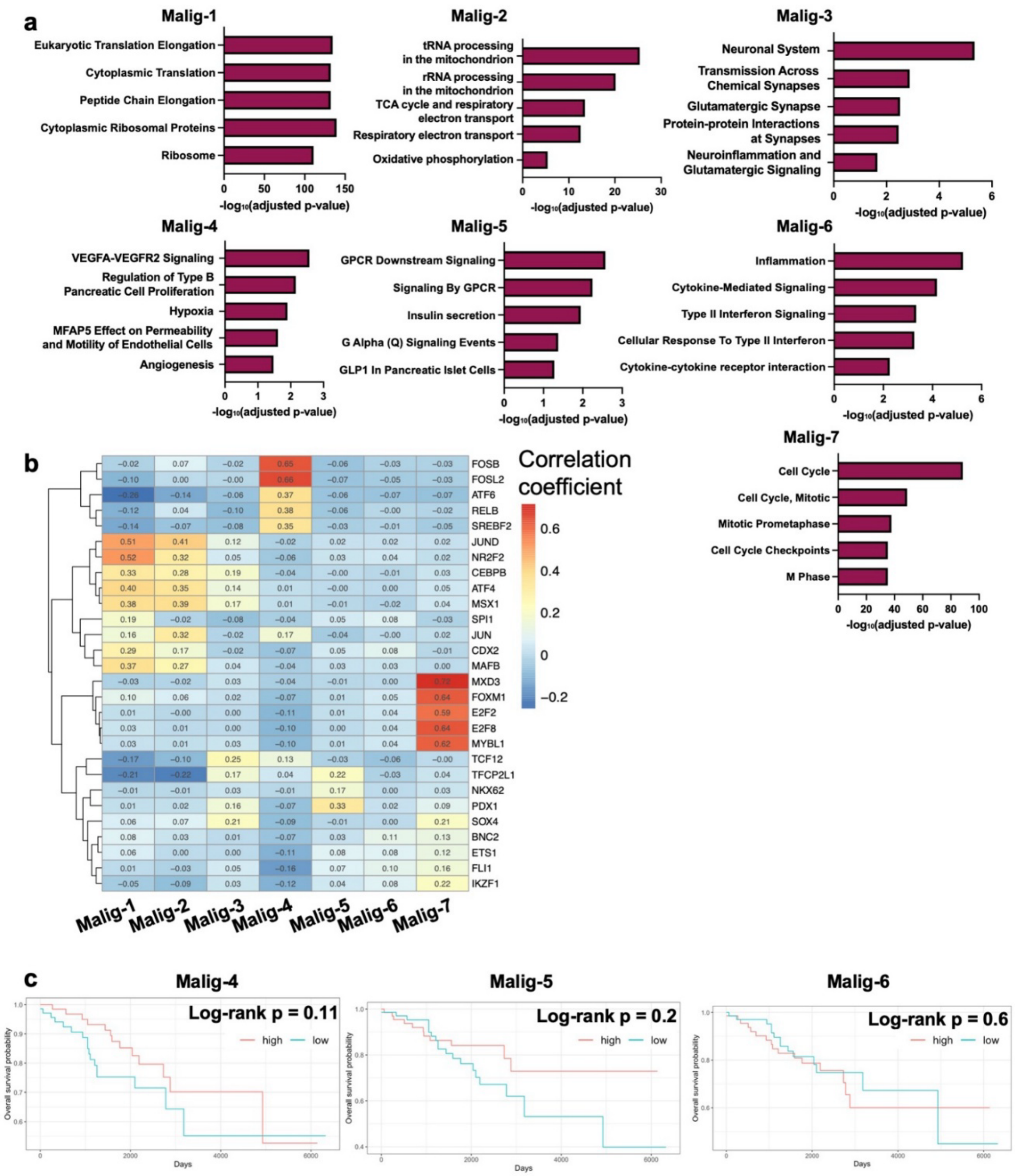


**Figure S4: Enriched gene ontology terms and gene regulatory networks in malignant GEPs across nonfunctional PNETs. (a)** Barplots of five selected terms from the top ten significant gene sets from EnrichR analyses (cutoff of adjusted p-value ≤ 0.05). Bars represent the -$\log_{10}$(Benjamini-Hochberg adjusted p-value). **(b)** The Pearson correlation coefficient was calculated between the SCENIC-derived score of transcription factor activity and the cNMF-derived GEP score. Heatmap shows the top 5 transcription factors with the highest, significant correlation coefficients for each GEP (columns) and hierarchical clustering was performed on the Pearson correlation coefficients (legend). **(c)** Kaplan Meier plots of overall survival stratified by high and low malignant GEP scores for Malig-4, Malig-5, and Malig-6.

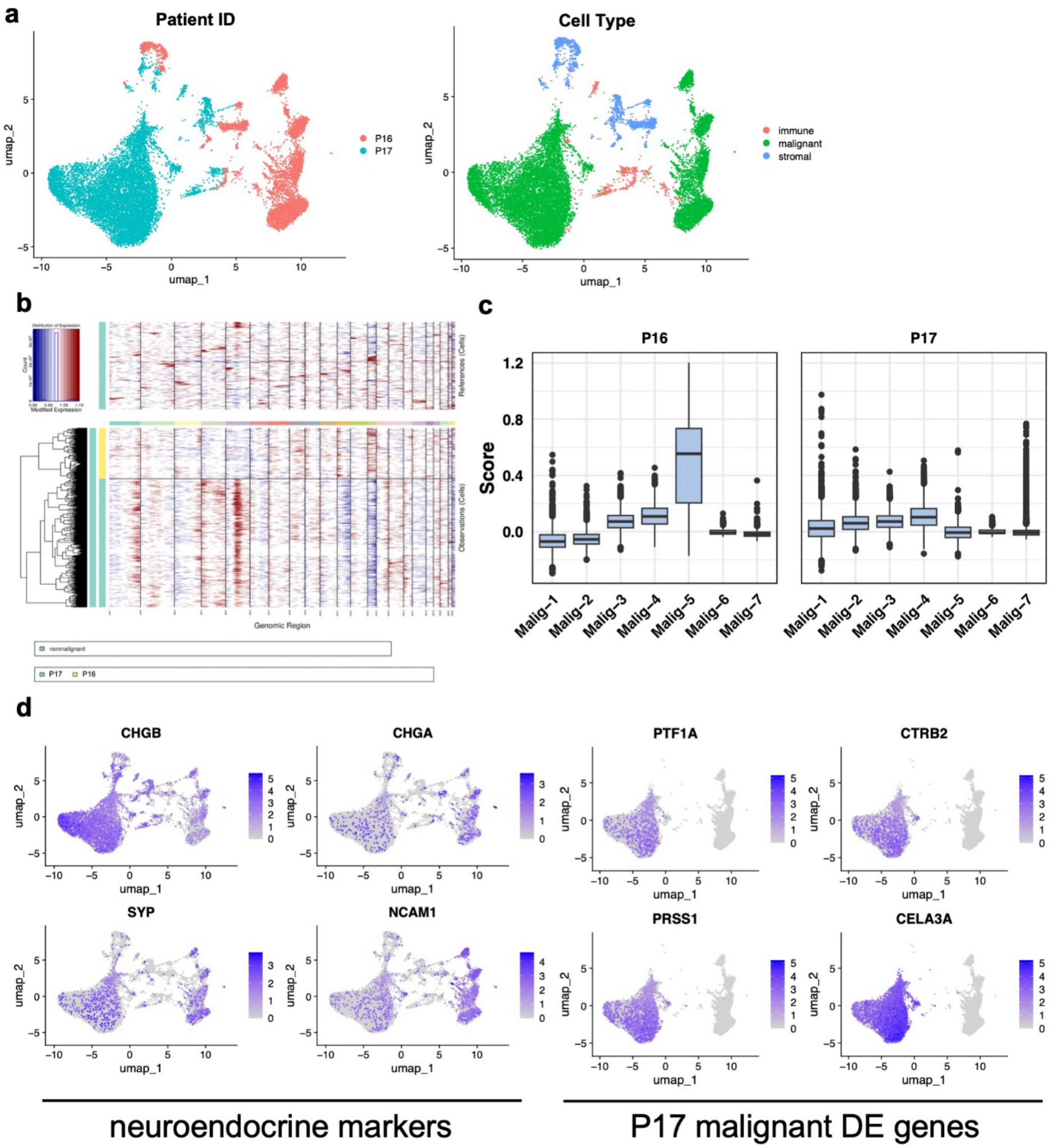


**Figure S5. Heterogeneity of malignant cell states in a case study of two PNETs with neoadjuvant treatment. (a)** UMAP embedding of single-nucleus profiles (dots) colored by sample (left) and cell type compartment (right). **(b)** Inferred amplifications (red) and deletions (blue) based on expression of a sliding 100-gene window in each chromosomal locus (columns) for each cell (rows). Green bar represents malignant cells from P16 and yellow represents malignant cells from P17. Top panel represents nonmalignant cells. **(c)** Boxplots of the score of each malignant GEP in the single nucleus profiles of the malignant cells in P16 and P17 colored by sample (legend). **(d)** Feature plots showing expression of neuroendocrine markers (left) and differentially expressed genes from P17 (right).

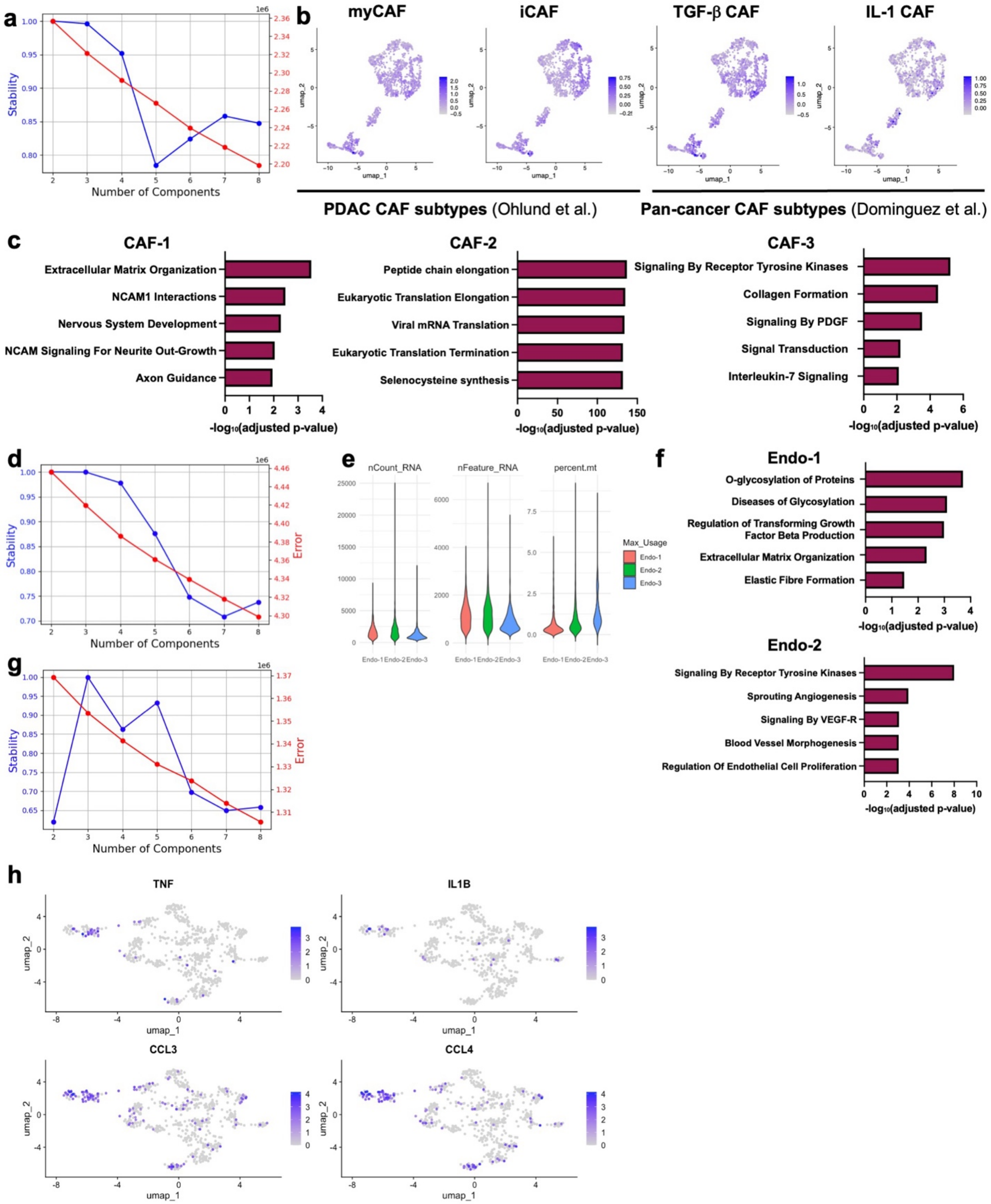


**Figure S6. Annotation of nonmalignant cell states in primary, nonfunctional PNET. (a)** Estimated stability (blue, left y-axis) and error (red, right y-axis) in the cNMF solution (left) learned with different programs (components, x-axis) for CAFs. **(b)** UMAP embedding of single nucleus profiles (dots) of fibroblast cells from primary, nonfunctional, untreated PNET tumors colored by the score (blue) of myCAF, iCAF, TGF-b CAF, and IL-1 CAF gene signatures. **(c)** Barplots of five selected terms from the top ten significant gene sets from EnrichR analyses for each CAF subtype (cutoff of adjusted p-value ≤ 0.05). Bars represent the $-\log_{10}$(Benjamini-Hochberg adjusted p-value). **(d)** Estimated stability (blue, left y-axis) and error (red, right y-axis) in the cNMF solution (left) learned with different programs (components, x-axis) for vascular endothelial cells. **(e)** Violin plots of number of features, number of counts, and percent mitochondrial genes for each endothelial cell GEP. **(f)** Barplots of top ten significant gene sets from EnrichR analyses for each endothelial subtype (cutoff of adjusted p-value ≤ 0.05). Bars represent the $-\log_{10}$(Benjamini-Hochberg adjusted p-value). **(g)** Estimated stability (blue, left y-axis) and error (red, right y-axis) in the cNMF solution (left) learned with different programs (components, x-axis) for macrophages. **(h)** Feature plots of various inflammatory genes in macrophages shown by level of expression (blue) on UMAP embedding of macrophages.

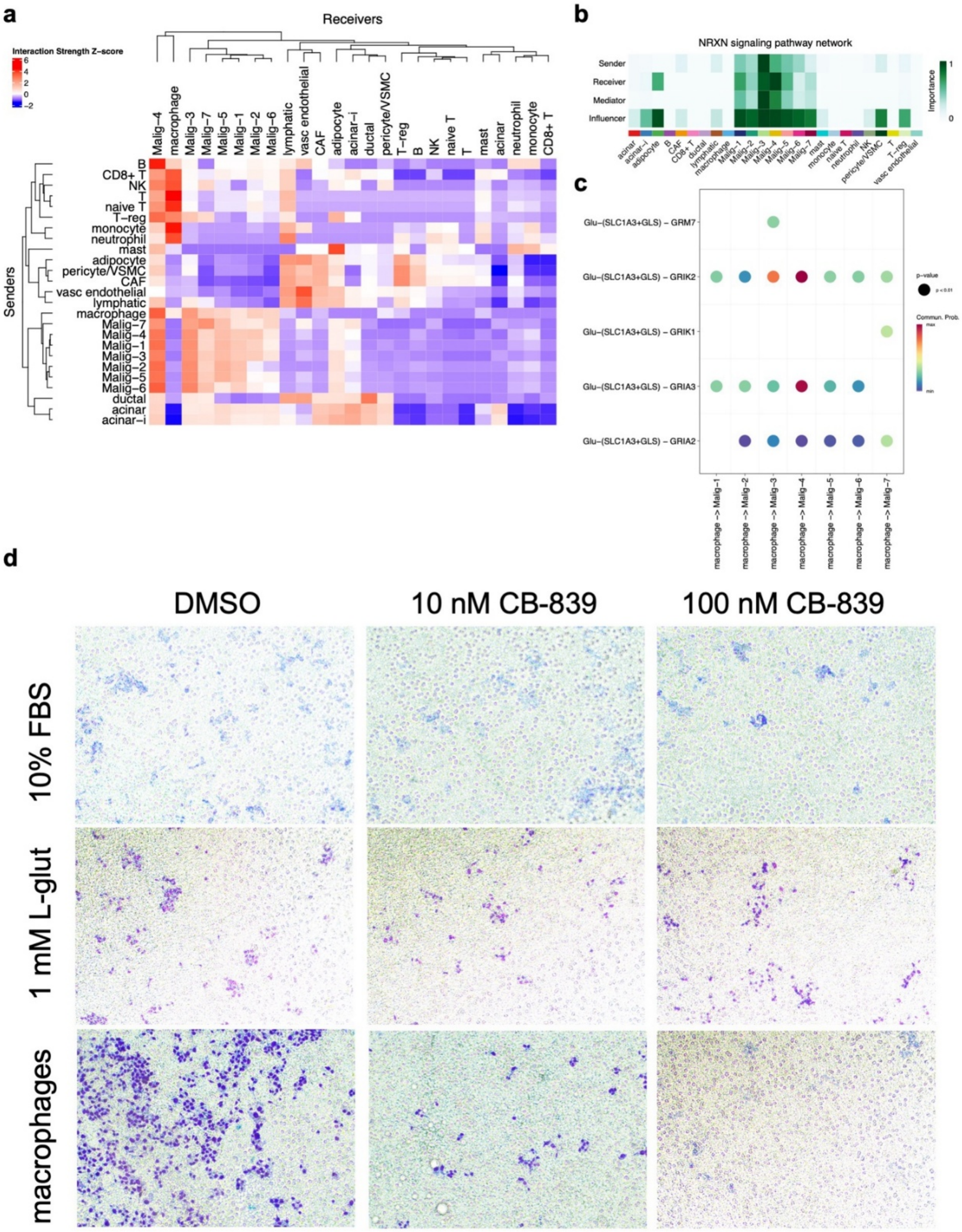


**Figure S7. Intercellular interactions in the PNET tumor microenvironment. (a)** Heatmap of interaction strength z-score (legend) between sender (rows) and receiver (columns) cell types inferred from single-nucleus profiles in primary, nonfunctional PNET. **(b)** Importance of different cell types (green) in the signaling of neurexins in the tumor microenvironment, with Malig-3 cells as the most important senders (top row) and other malignant subtypes as the most important receivers (second row). **(c)** Bubble plot showing significant ($p < 0.01$) ligand-receptor pairs involved in glutamate signaling between macrophages and malignant GEPs. Color bar represents relative probability. **(d)** Representative images of transwell migration assay in Figure 4f.

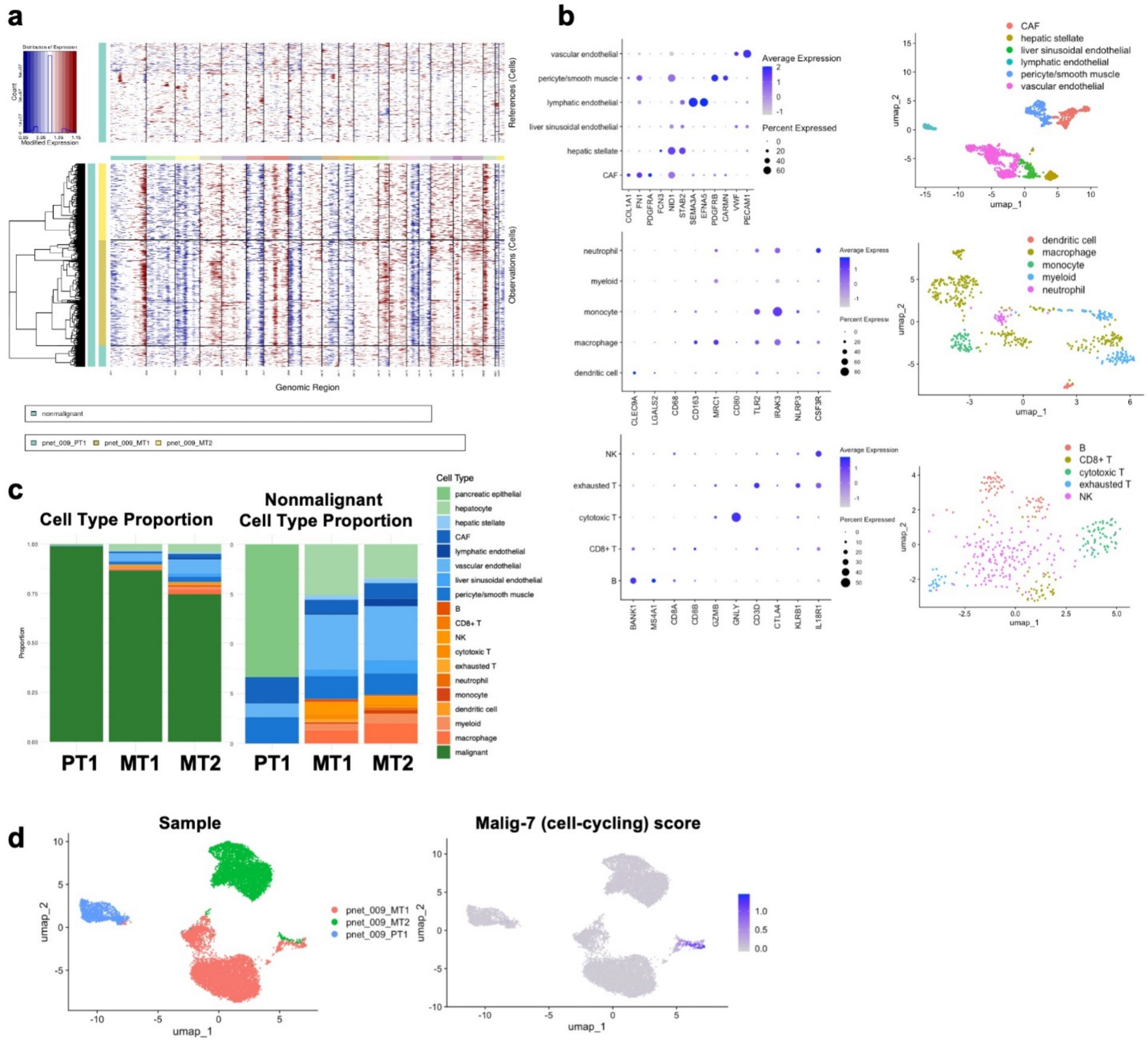


**Figure S8. Transcriptomic landscape of a case study of a nonfunctional primary PNET with two asynchronous hepatic metastases. (a)** Inferred amplifications (red) and deletions (blue) based on expression of a sliding 100-gene window in each chromosomal locus (columns) for each malignant cell of P18 (rows) from sample PT1 (turquoise bar), MT1 (ochre bar), and MT2 (yellow bar). **(b)** Average expression (color) of selected marker genes (columns) across annotated cell types (rows) in the stromal (top), myeloid (middle), and lymphoid (bottom) compartments with corresponding UMAP embeddings colored by cell type (right). **(c)** Stacked barplot of cell type proportion of each sample of P18 including malignant cells (left) and of only nonmalignant cells (right). **(d)** UMAP of malignant cells from P18 colored by sample (top) and Cell-cycling GEP score (bottom).
